# Identification of conserved gene regulatory networks that integrate environmental sensing and growth in the root cambium

**DOI:** 10.1101/839464

**Authors:** Goh Choe, Nam Van Hoang, Yi Zheng, Ana Cecilia Aliaga Fandiño, Jaeryung Hur, Inyoung Sung, Hongryul Ahn, Sun Kim, Zhangjun Fei, Ji-Young Lee

## Abstract

Cambium drives lateral growth of stems and roots, contributing to diverse plant growth forms. Root crop is one outstanding example of the cambium-driven growth. To understand its molecular basis, we used radish to generate a compendium of root tissue- and stage-specific transcriptomes from two contrasting inbred lines in root growth. Expression patterns of key cambium regulators and hormone signaling components were validated. Clustering and GO enrichment analyses of radish datasets followed by comparative analysis against the newly established Arabidopsis early cambium data revealed evolutionary conserved stress-response transcription factors that might intimately control the cambium. Indeed, *in vivo* network made of selected stress-response and cambium regulators indicated *ERF-1* as a potential key checkpoint of cambial activities, explaining how the cambium-driven growth is altered in response to environmental changes. Together, this study provides rich information about dynamic gene expression changes along the cambium-driven root growth with future engineering schemes for crop yields.

## Introduction

Plant growth throughout life cycles is driven by meristems, reservoirs of stem cells and their daughters, which actively divide to generate cells constituting organs. Cambium is a meristem that is established during postembryonic development. The cambium promotes the growth of stems and roots in the radial direction by generating cells of xylem tissue toward the organ center and phloem tissue toward the organ periphery. This secondary growth is critical for the biomass increase in most of the perennial tree species. Despite the importance of secondary growth, its underlying physiological and molecular networks have been just begun to be identified through the studies of Arabidopsis (Miyashima et al., 2013; Zhang et al., 2011; Zhang et al., 2014).

Plant roots absorb water and nutrients from the soil and transport them to the rest of plant bodies through the vascular system. As part of this activity and to support the above-ground body parts, roots continuously grow and branch out making a complex root system. In addition, several biennial and perennial plants use roots for storage. These include carrots, sweet potatoes, and cassavas, which are cultivated as very important food sources.

Root crops originate from the thickening of primary or lateral roots (Tonn and Greb, 2017). Thickening process of roots is mainly mediated by cambium. Considering that root crops serve as a main staple food in environmentally marginal regions, we need to understand the cambium regulatory programs underlying storage root development and to develop strategies making root crops with sustainable yields. Radish, *Raphanus sativus* L. cultivated in eastern Asia, develops a very large storage tap root weighing more than 2kg in just two months of growth in the field. We previously reported that cell division activity in the cambium determines radish root yields (Jang et al., 2015). Radish can serve as one of the ideal root crop models for the following reasons. First, the radish draft genome is available (Jeong et al., 2016; Kitashiba et al., 2014). Second, the growth time for its storage roots is much shorter than other root crops. Third, unlike other root crops, radish root does not require environmental stimuli for storage root formation. Lastly, the evolutionary proximity of radish to Arabidopsis makes comparative analysis feasible.

In this study, we developed tissue-specific transcriptome of radish storage tap roots in two inbred lines with contrasting storage root growth and yields. These data allowed us to identify novel regulators of cambium activities and many non-coding RNAs that might tune the expression of protein-coding genes. Through comparative analyses with the cell-type-specific gene expression data in Arabidopsis roots (Zhang et al., 2019), we identified co-expression gene regulatory networks (GRNs) conserved in Arabidopsis and radish roots. These included a GRN highly enriched with transcription factors (TFs) involved in biotic/abiotic stress responses. The inference of *in vivo* GRN amongst selected TFs using knockout mutants in Arabidopsis roots predicted that ERF-1 might serve as one of the key nodes of the network. Interestingly, both increasing and decreasing ERF-1 levels negatively affected the root secondary growth, suggesting that ERF-1 might function as a balancer. Our study newly finds stress response regulator(s) as a critical checkpoint for the cambium mediated growth and offers a direction to biomass engineering for controlling yields.

## Results and Discussion

### Tissue-specific profiles of protein-coding and non-coding RNAs in radish storage tap roots

We generated tissue-specific transcriptome profiles in the radish tap root using Laser Capture Microdissection (LCM) coupled with RNA sequencing (RNA-Seq). We selected two inbred lines, 216 and 218, which exhibit contrasting tap root growth and yields (Jang et al., 2015). We sampled their roots at three growth stages, 5, 7, and 9 weeks post seed planting, based on radish root growth dynamics (Supplementary Fig. 1b). From root sections, cambium (Ca) and tissues adjacent to cambium, which are mainly composed of parenchyma cells, named as cortex for phloem side (Co) and parenchyma for xylem side (Pa), were cut out and collected with laser, and then processed for paired-end RNA-Seq library construction (Supplementary Fig. 1c; Fig. 1a).

**Figure 1.**
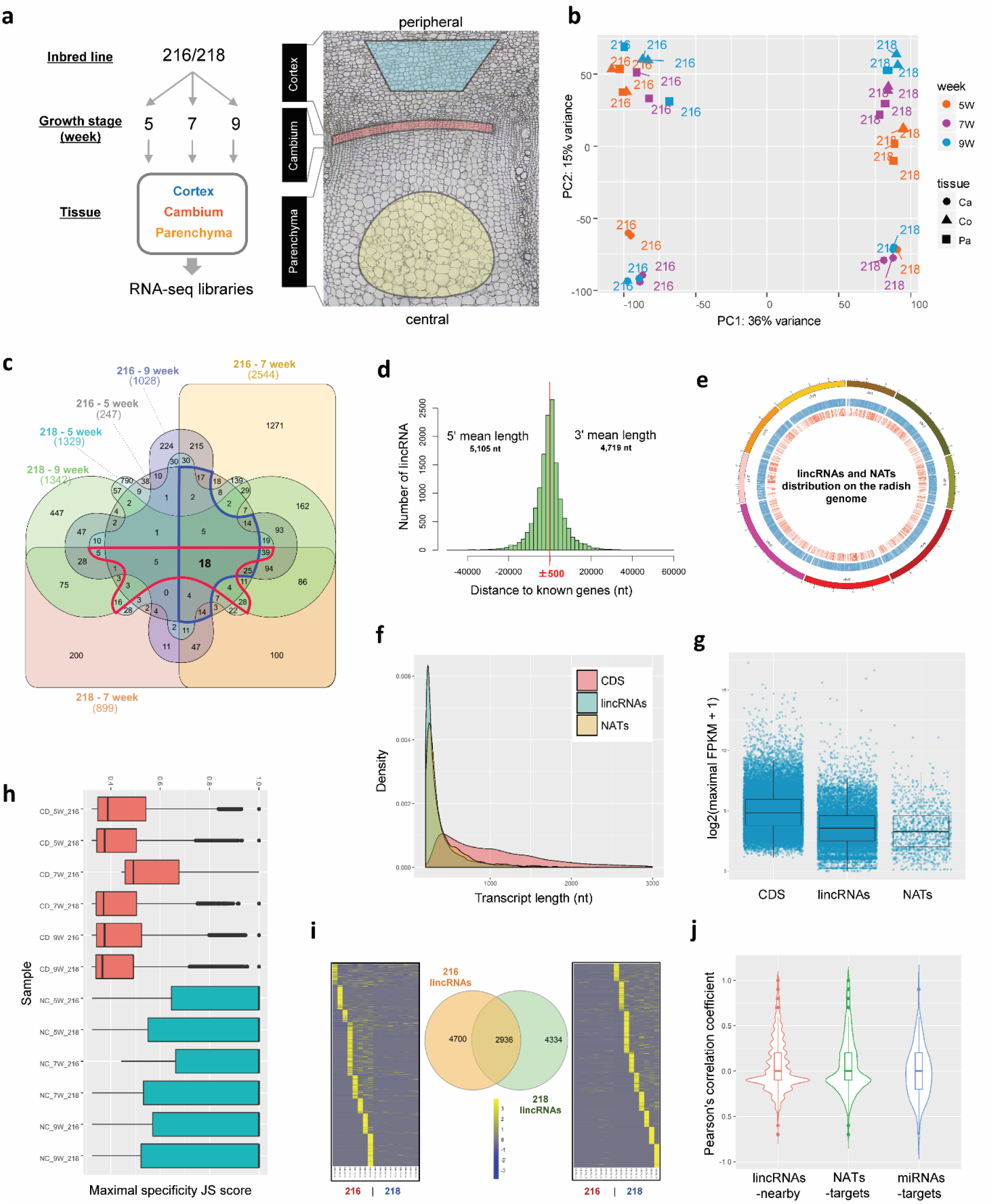
RNA-Seq experiment design, differentially expressed gene identification, and lincRNA analysis. **a** RNA-Seq library sample preparation design (left) and representative example of tissue areas for laser capture microdissection (right). Region of target areas for sampling was marked and colored in blue, red, and yellow for cortex, cambium, and parenchyma respectively. **b** Principal component analysis of RNA-Seq expression data. Top five thousand genes based on their variance rank were selected as informative genes, and used for PCA analysis. The number of samples for each stage is indicated in the legend, with shape indicating tissue type (Ca, cambium; Co, cortex; and Pa, parenchyma) and color indicating growth stage (5W, 5 weeks; 7W, 7 weeks; and 9W, 9 weeks). **c** Venn diagram showing DEG analysis (|fold change| >2 and FDR <0.05). Cambium DEGs in 216 and 218 were compared to show overlapping genes and uniquely expressed genes between the two inbred lines. Blue and red lines indicate 99 and 118 DEGs that were shared throughout the growth stages in lines 216 and 218, respectively. **d** Distance of lincRNAs to known protein-coding genes. **e** Circos plot of distribution of lincRNAs and natural antisense transcripts (NATs) on the radish genome. Outer lines represent nine radish chromosomes, the blue bars (middle) represent lincRNA distribution, and orange bars (inner) represent NAT distribution. Only lincRNAs that were assigned to nine chromosome pseudomolecules of the radish genome version Rs 1.0 (Jeong et al., 2016) were shown. **f** Length distribution of identified lincRNAs and NATs in comparison with protein-coding transcripts. **g** Expression level comparison between lincRNAs and protein-coding transcripts. The maximal expression level of each transcript across samples was used. Data was log2 transformed for visualization. **h** The distribution of maximal tissue specificity score (JS score) of lincRNAs and coding transcripts, calculated for each stage (5 weeks, 7 weeks and 9 weeks) of cambium, cortex and parenchyma tissues from line 216 and 218. CD denotes coding transcripts, while NC denotes lincRNAs. **i** Venn diagram of tissue-specific transcripts unique to line 216 and 218 in each stage and each tissue. **j** Pearson correlation coefficient between lincRNAs-nearest genes, NATs-target genes, and miRNAs-target genes, respectively.

We generated a total of ~920 million raw read pairs of RNA-Seq data. After removing low quality and contaminated sequences, we obtained a total of ~607 million high-quality cleaned read pairs from 34 RNA-Seq libraries, with each library having 6-70 million read pairs (Supplementary Table 1A). Approximately 70% of high-quality cleaned reads were mapped to the radish reference genome (http://radish-genome.org/). In addition, we performed *de novo* assembly of these high-quality cleaned reads and identified a total of 3,515 novel protein-coding genes that are not present or annotated in the reference genome. These novel genes were combined with the reference genes for the downstream gene expression analysis (Supplementary Fig. 2; Supplementary Tables 1B-D).

A principal component analysis (PCA) for protein-coding transcripts using the regularized log-transformed expression values of 5,000 most variable genes across the samples (Fig. 1b) identified one outlier from Co in 7-week-old roots of line 216. In the PCA with the remaining 33 RNA-Seq libraries after removing this outlier, the first principal component clearly separated the two inbred lines, accounting for 36% of the total variance. In each inbred line, Ca samples grouped distinctively from Co and Pa along the second principal component. This can be explained by the fact that both Co and Pa are mostly parenchyma cells though they are physically separated by cambium. Ca samples of line 216 grouped apart from the early and late growth stages, while all the Ca samples of 218 grouped closely regardless of the growth stages.

To identify differentially expressed genes (DEGs), we conducted pairwise comparisons of the Ca against Pa and Co in each growth stage (|fold change| >2 and false discovery rate (FDR) <0.05; Supplementary Tables 2A and B). Among a total of 4,602 DEGs, 3,406 genes were differentially expressed in only one of the stages in either 216 or 218. Remaining 1,196 DEGs were shared between the two lines. 99 DEGs were shared throughout the growth stages in line 216, and 118 were in line 218 (Fig. 1c). Only 18 DEGs were shared between the two (Fig. 1c; Supplementary Fig. 3). These data collectively suggest the dynamic gene expression changes during the storage root growth.

Large intergenic non-coding RNAs (lincRNAs) could play critical roles in various biological processes including transcriptional and post-transcriptional regulations of protein-coding genes in both animals and plants (Liu et al., 2015; Ulitsky and Bartel David, 2013). We identified a total of 14,202 lincRNAs that were within 500 nucleotides either upstream or downstream of the nearest protein-coding genes (Supplementary Tables 3A-C; Fig. 1d, e). Consistent with several reports (Golicz et al., 2018; Wang et al., 2015), lincRNAs identified in radish were shorter in length (Fig. 1f) and had lower expression levels than those of protein-coding transcripts (Fig. 1g). We further validated the expression of 32 out of 40 selected lincRNAs using reverse transcription PCR (RT-PCR) of RNAs extracted from 7-week radish roots from lines 216 and 218 (Supplemental Table 9A; Supplementary Fig. 4).

We further compared the tissue specificity of lincRNAs and coding transcripts (Fig. 1h). Distinct distribution patterns of maximal tissue specificity score [Jensen-Shannon (JS) score (Cabili et al., 2011)] of lincRNAs and coding transcripts indicated that lincRNAs tend to be more tissues-specific than coding transcripts. A similar trend was also reported in human (Cabili et al., 2011), cotton (Wang et al., 2015) and soybean (Golicz et al., 2018). With JS score above 0.6, 4,700 and 4,334 lincRNAs in lines 216 and 218 were found to be preferentially expressed in the Ca, Co and Pa at different stages (Fig. 1i). We then further asked the potential roles of lincRNAs in the expression of coding transcripts. First, we analyzed the correlative expression of lincRNAs and nearest protein-coding transcripts. Among 7,732 pairs subject to analysis (Fig. 1j; Supplementary Table 4A; Supplementary Fig. 5), 1,533 pairs of lincRNA-coding transcript exhibited positive correlation (*r* ≥ 0.3) whereas only 397 pairs showed negative correlation (*r* ≤ −0.3). This is in agreement with previous studies (Ding et al., 2018) in which more lincRNAs have positive regulatory influence on the transcription of their nearby mRNAs.

Second, we identified 2,237 *trans* natural antisense transcripts (*trans-*NATs, ~16% total lincRNAs) that potentially target 3,270 protein-coding transcripts (Fig. 1e, f). The expression of eight selected NATs (LIN17 - LIN25) showing evolutionary conservation against Arabidopsis TAIR10 intergenic sequences was validated in RT-PCR (Supplementary Fig. 4). Pearson’s correlation analysis between *trans*-NATs and their potential target protein-coding transcripts identified 194 pairs with positive correlation (*r* ≥ 0.3) and 68 pairs with negative correlation (*r* ≤ −0.3) (Fig. 1j; Supplementary Table 4B; Supplementary Fig. 6). On a genome-wide scale, the expression of NATs was generally associated with the down-regulation of their target sense genes through several mechanisms resulting in the suppression of target gene expression at the transcriptional level by histone modification or chromatin modifiers, or by post-transcriptional regulation (Liu et al., 2010; Ponting et al., 2009). However, there are studies indicating that NATs are associated with transcriptional activation, or up-regulation of the target genes (Jabnoune et al., 2013; Li et al., 2008; Xu et al., 2017). Unlike *cis*-NATs which are known to be transcribed from ~30% of protein-coding genes in Arabidopsis (Yamada et al., 2003), and were more extensively studied, *trans*-NATs are derived from different loci and only a few have been studied. In that context, our results provide a platform for understanding how trans-NATs affect their target coding transcripts in a positive or negative manner.

Lastly, a total of 154 lincRNAs were found to match 72 miRNA families in the mirBase release 22 (Kozomara and Griffiths-Jones, 2014). Among these, 32 lincRNAs corresponding to 12 miRNA families were retained after confirmation of their secondary structure (see Supplementary Table 4C for a list of 32 precursor miRNA sequences and the associated lincRNAs; Supplementary Fig. 7 for their secondary structure). Using predicted mature miRNAs (20-22 nt) from these precursor miRNAs, we identified 189 pairs of miRNA-target gene. As indicated in several studies (Gan and Denecke, 2013; Lee et al., 2008), the expression of mature miRNAs is highly concordant with that of their respective precursor miRNAs. Thus, we used the sum of expression values from the miRNA-encoding lincRNAs and target coding genes in the Pearson’s correlation analysis. Among 189 pairs, we detected 26 pairs showing negative correlation (*r* ≤ −0.3; Supplementary Table 4D and Supplementary Fig. 8).

### Validation of the regulators of vascular tissue development in the tissue-specific transcriptome data

We validated the laser-captured RNA-Seq data by examining the expression of putative orthologues of genes already known to control the vascular tissue development (Fig. 2a). *PXY* (*PHLOEM INTERCALATED WITH XYLEM*) encoding a receptor-like kinase promotes cambium cell proliferation through *WOX4* (*WUSCHEL-RELATED HOMEOBOX 4*) and *WOX14* (Etchells et al., 2013). In our RNA-Seq data, expression levels of radish orthologs of these genes were higher in the Ca than in neighboring tissues in all the stages, except for *RsWOX14* (*Raphanus sativus WOX14*) in 9-week-old roots of line 216. Radish orthologs of *MOL1* (*MORE LATERAL GROWTH 1*), another receptor-like kinase regulating cambium activity, also showed high expression in the cambia of both inbred lines (Agusti et al., 2011). *AGL16* (*AGAMOUS-like 16*), a MADS-box family TF reported to promote the secondary growth in the sweet potato root (Noh et al., 2010), also showed enriched expression in the Ca.

**Figure 2.**
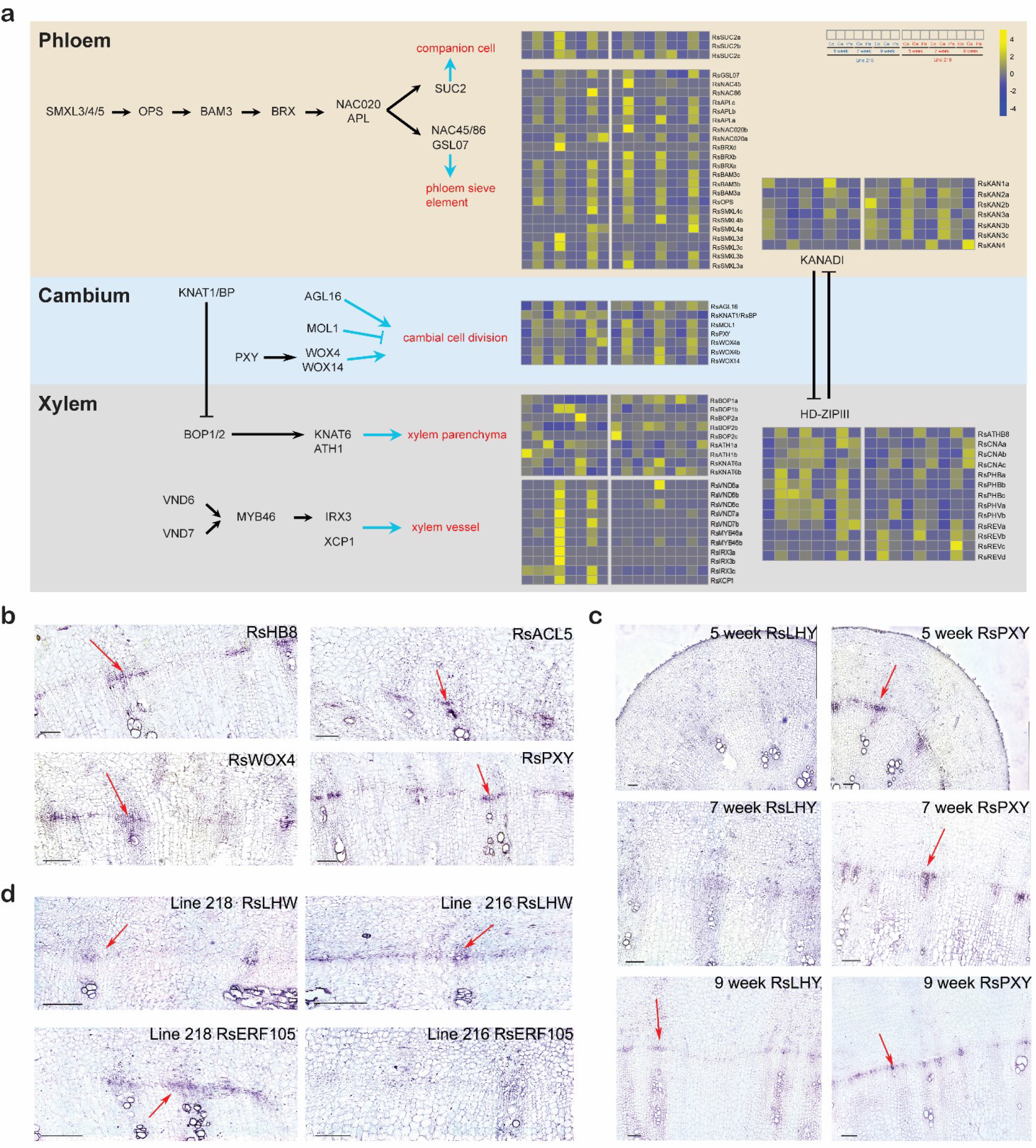
Expression patterns of radish gene homologs of Arabidopsis genes known for vascular tissue patterning and differentiation. **a** Schematic representation of vascular tissue patterning in Arabidopsis was shown with heatmaps visualizing of the expressions of radish homologs. Gene-wise normalized FPKM values were used for the heatmaps. The order of sample was shown on the top right. Co, cortex; Ca, cambium, and Pa, parenchyma. **b-d** show *in situ* hybridization antisense probe results. **b** shows cambium enriched genes: *RsHB8*, *RsACL5*, *RsWOX4* and *RsPXY* in 7-week radishes of line 216. Confined signal (purple) appeared in the cambium cells. **c** shows a time course RNA-Seq validation in line 216 radishes of 5, 7 and 9 weeks. Image shows the different expression dynamics between *RsLHY* and *RsPXY*. **d** shows cross-line comparison validation in 7-week radishes of line 216 and line 218. On the top, *RsERF105*, only expressed in line 218 and on the bottom *RsLHW* expressed significantly more in line 216 than in line 218. Magnification 100X, 200μm scale bar.

Genes associated with the development of phloem (Anne and Hardtke, 2018; Furuta et al., 2014), such as radish homologs of *SMXLs (SUPPRESSOR OF MAX2 1-LIKE 3,4, and 5*), *OPS* (*OCTOPUS*), *BAM3* (*BARELY ANY MERISTEM3*), and *BRX* (*BREVIS RADIX*) involved in phloem development and differentiation; *NAC20* (*NAM, ATAF1,2 and CUC2 20*) and *APL* (*ALTERED PHLOEM DEVELOPMENT*) controlling sieve element enucleation process through *NAC86/45*; *GSL07* (*GLUCAN SYNTHASE-LIKE 07*) encoding a callose synthase responsible for phloem maturation; and *SUC2* (*SUCROSE-PROTON SYMPORTER 2*) mediating phloem loading processes in companion cells were mostly enriched in the Ca samples. Xylem regulators, *VND* (*VASCULAR-RELATED NAC DOMAIN*) genes, *MYB46* (*MYELOBLASTOSIS 46*) regulating cellulose synthase gene, and *IRX3* (*IRREGULAR XYLEM 3*) and *XCP1* (*XYLEM CYSTEINE PEPTIDASE 1*) responsible for the autolysis of xylem vessels (Furuta et al., 2014; Schuetz et al., 2013), were also enriched in the Ca samples. Noticeably, the expression of xylem regulators was induced in 7- and 9-week-old roots of the 216 line that undergo active growth. *BP*/*KNAT1* (*BREVIPEDICELLUS*/*KNOTTED-LIKE FROM ARABIDOPSIS THALIANA 1*), expressed in the cambium and xylem vessels of Arabidopsis roots, was reported to prevent *BOPs* (*BLADE ON PETIOLEs*) from expanding to the xylem domain (Liebsch et al., 2014; Woerlen et al., 2017). In the Arabidopsis *bp* mutant, *BOPs* promoted the formation of xylem parenchyma by expanding to the xylem side. Interestingly, our radish data indicated the broad expression of *RsBOPs* including parenchyma of the xylem side. In addition, *RsKNAT6*s and *RsATH1*s, which are downstream of *BOPs*, showed expression patterns similar to *BOP*. These data suggest that in radish the KNAT1-BOP pathway might have been modified in a direction that promotes the fleshy parenchyma formation instead of xylem vessels.

HD-ZIP III (Class III homeodomain leucine zipper) and KANADI gene families are important for establishing organ polarity in Arabidopsis (Emery et al., 2003). In vascular tissues, HD-ZIP III’s are enriched in the xylem side and KANADIs are in the phloem side to antagonistically regulate each other’s activity and expression domains (Emery et al., 2003). Consistent with this, *RsKANADI*s were expressed highly in the Co, and *RsCNA*s (*RsCORONA*), *RsPHB*s (*RsPHABULOSA*) and *RsPHV*s (*RsPHAVOLUTA*) showed strong expression in the Pa as well as in the Ca. Interestingly, five HD-ZIP IIIs (two *PHAVOLUTA*, two *REVOLUTA*, one *PHABULOSA*), which are involved in the boundary between xylem and phloem tissues, negatively correlated with the expression of *Rsa-miR*166. *Rsa-miR*166 showed high enrichment in the Co, which seems to actively exclude *HD-ZIP III*s in this region (Supplementary Table 4D).

For visual validation, we selected five genes and examined their expression patterns using RNA *in situ* hybridization (Supplementary Fig. 9; Supplementary Tables 9B-D). *RsACL5*, *RsATHB8*, *RsPXY*, and *RsWOX4* analyzed in 7-week-old roots of line 216 indicated their specific expression in subsets of cambia (Fig. 2b; Supplementary Fig. 10a). *RsPXY* was further examined in time course in line 216 together with *RsLHY*, which has not been described in relation to cambium. *RsPXY* showed cambium specific expression throughout the selected stages whereas *RsLHY* did only in the 9-week-old root of line 216, precisely reflecting their expression patterns based on RNA-Seq data (Fig. 2c; Supplementary Fig. 9 and 10b). These indicate that our RNA-Seq data accurately reflect the *in vivo* transcriptome profiles. In addition, our RNA *in situ* hybridization revealed that expression of genes involved in the cambial identity and maintenance and those regulating cell-type specification are enriched in only a few layers of cambial zones. This suggests that the expression of these regulators is dynamically turned on and off in the cambial zone while the cambium rapidly produces cells for growth.

### Evidence for cambial activities via integration of auxin-cytokinin signaling

Plant hormones are important internal regulators of the secondary growth. Auxin and its signaling processes have been known to promote vascular patterning through positive feedback regulation (Miyashima et al., 2013). The concentration gradient of auxin was reported to correlate with cambial activity (Campbell and Turner, 2017; Uggla et al., 1998). To examine the influence of the auxin pathway, we analyzed DEGs associated with auxin biosynthesis and signaling (Supplementary Fig. 11). In the cambium, two *RsAAOs* (*indole aldehyde oxidase*) and one *RsYUC6* (*YUCCA6*), which catalyze the final steps of the auxin biosynthesis, and *RsGH3.6* genes (*GRETCHEN HAGEN 3.6*), which encodes an auxin conjugating enzyme, showed enriched expression. This suggests that *de novo* auxin biosynthesis and subsequent modification occur actively in the cambium (Supplementary Fig. 11a). Expression patterns of auxin transporters were also in line with the high demand of auxin in the cambium (Supplementary Fig. 11b). These transporters included plasma membrane localized transporters mediating cell-to-cell auxin transport, and organelle localized transporters controlling auxin homeostasis. Among *ARFs*, *RsARF5/MP* associated with transcriptional activation of many auxin-responsive genes were preferentially expressed in the cambium, which was consistent with the expression patterns shown in Arabidopsis stems (Brackmann et al., 2018). *RsARF10* and *RsARF16*, transcriptional repressors, showed relatively weak expression in the cambium. A subset of *AUX/IAAs* known as auxin-responsive genes (Abel and Theologis, 1996; Paponov et al., 2008) were also enriched in the cambium.

Cytokinin plays an important role in regulating cambium activity (Matsumoto-Kitano et al., 2008; Nieminen et al., 2008). When we checked the differential expression of cytokinin biosynthesis and signaling genes (Supplementary Fig. 12), we found four *RsIPTs* (*ISOPENTENYLTRANSFERASE*) highly enriched in cambium. A recent study using Arabidopsis embryo underscored the importance of cytokinin transporters in activating cytokinin signaling (Zurcher et al., 2016). In that study, the presence of cytokinin importer in the plasma membrane negatively influenced cytokinin signaling, suggesting that cytokinin needs to be exported to be perceived by plasma membrane localized receptor proteins. In our RNA-Seq data, cytokinin exporters were enriched in cambial tissues, whereas cytokinin importers were depleted in cambial tissues, which is in line with the former hypothesis. A-type ARR (ARABIDOPSIS RESPONSE REGULATOR) genes and *RsAHPs* (*ARABIDOPSIS HISTIDINE PHOSPHOTRANSFER PROTEINS*) that are cytokinin responsive (Muller and Sheen, 2007) were enriched in the cambium, indicating the high cytokinin level in the cambium. In contrast to the auxin and cytokinin pathway components that were extensively represented in the DEG list, only a few associated with gibberellic acid and brassinosteroid pathways were found in the DEG list (Supplementary Fig. 13).

In Arabidopsis root procambium, auxin and cytokinin promote periclinal divisions by balancing the positive regulation mediated by *TMO5* and *LHW* and the negative regulation via thermospermine (Fig. 3a) (Vaughan-Hirsch et al., 2018). In our RNA-Seq data, genes involved in these pathways were differentially expressed. *RsTMO5* (*TARGETS OF MONOPTEROS*), *RsT5Ls* (*TMO5-LIKEs*), *RsLHW* (*LONESOME HIGHWAY*), and *RsLLs* (*LONESOME HIGHWAY LIKEs*), which promote periclinal cell divisions (Fig. 3b; Supplementary Fig. 1a), were enriched in the radish cambium. In particular, the expression of *RsLHW* was higher in line 216 than 218, which we further confirmed using RNA *in situ* hybridization (Fig. 2d). In the case of thermospermine pathway, which represses TMO5-LHW, thermospermine synthase *RsACL5* (*ACAULIS5*) was enriched in cambial tissues with higher expression in line 216. However, the expression of *RsSACL3*, an immediate inhibitor of TMO5-LHW, was weak in the cambium and enriched in the cortex. These expression dynamics collectively explain why the periclinal cell divisions are more active in line 216 than in the 218. Since involvement of *LHW* in the regulation of cambial activities has never been reported, we examined the secondary growth in the *lhw* mutant of Arabidopsis. As predicted, the *lhw* root showed a significant reduction in the secondary growth in comparison to the wild type (Fig. 3c).

**Figure 3.**
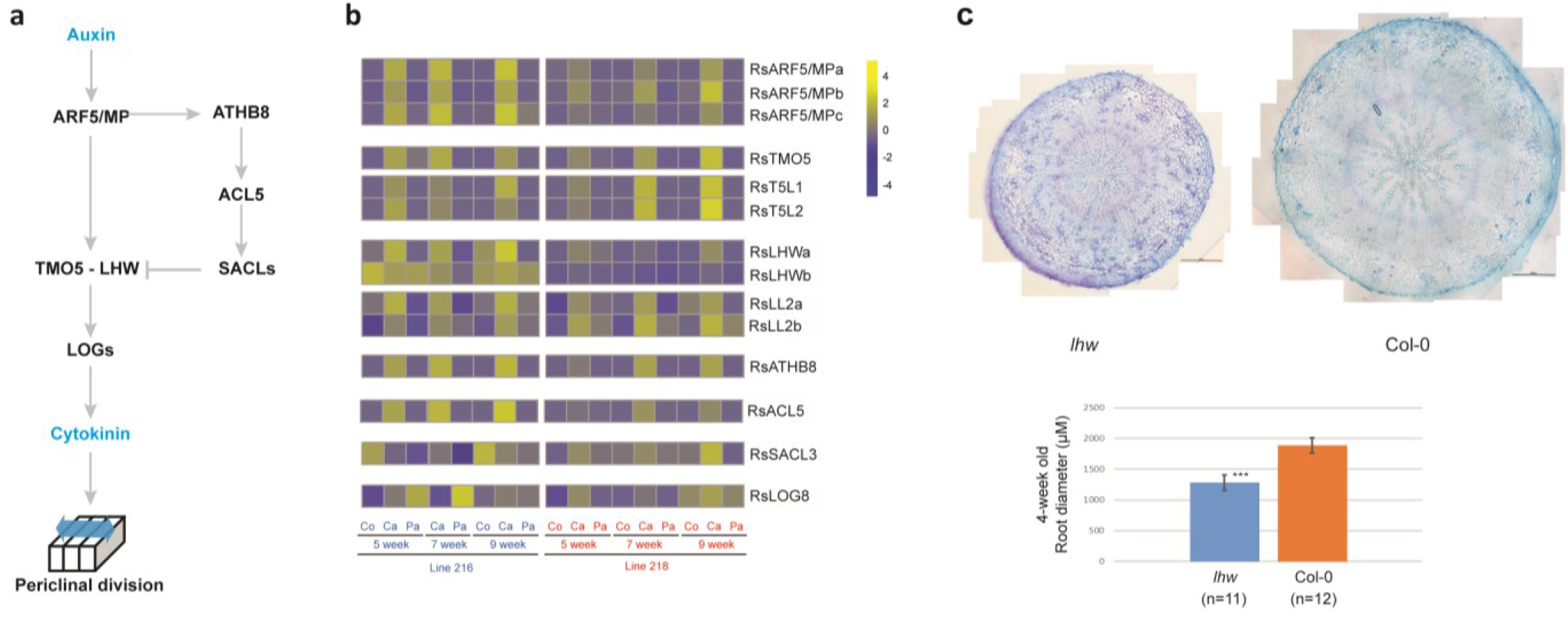
LHW pathway. **a** Schematic representation of known *LHW* pathway in Arabidopsis vascular development. **b** Expressions of radish homologs of Arabidopsis *LHW* pathway genes. Gene-wise normalized FPKM values were used for the heatmaps. The order of sample was shown at the bottom of the heatmap. Co, cortex; Ca, cambium, and Pa, parenchyma. **c** Representative cross section images of 4 week-old *lhw* and Col-0 roots (upper). Quantification of root cross sections of 4 week-old *lhw* and Col-0 root (lower). Asterisks; statistical significance based on Student’s t-test (*** p < 0.01). Scale bar, 200μm.

### Discovery of a stress response gene regulatory network conserved in Arabidopsis and radish root cambia

To comprehensively understand temporal and spatial gene expression dynamics during the storage root growth, we performed K-means clustering for a total of 4,602 DEGs obtained from the pairwise comparisons (Fig. 4a) (Lloyd, 1982). Among the 24 gene clusters, 20 showed preferential expression in one of the tissues in the root. Some of these also showed distinct expression dynamics between the two inbred lines. To gain insights into their functional associations, we conducted gene ontology (GO) analysis for each cluster (Fig. 4b; Supplementary Table 5). The enrichment of GO term associated with growth (GO:0040007) in K13 and 23, which showed cambium preferential expression in both 216 and 218, reflects the importance of growth-associated processes in the cambial cell activities. This term was also enriched in K14 and K20, which show much higher enrichment in the cambium of line 216 than in line 218, consistent with the faster growth in line 216 root. Contrary to K14 and K20, K5 showed much higher gene expression in the cambium of line 218, and its members were enriched with genes involved in response to endogenous and external stimuli. A similar trend was found for K1. These data indicate that the secondary growth-related process and the response to stimuli/stresses might be negatively related.

**Figure 4.**
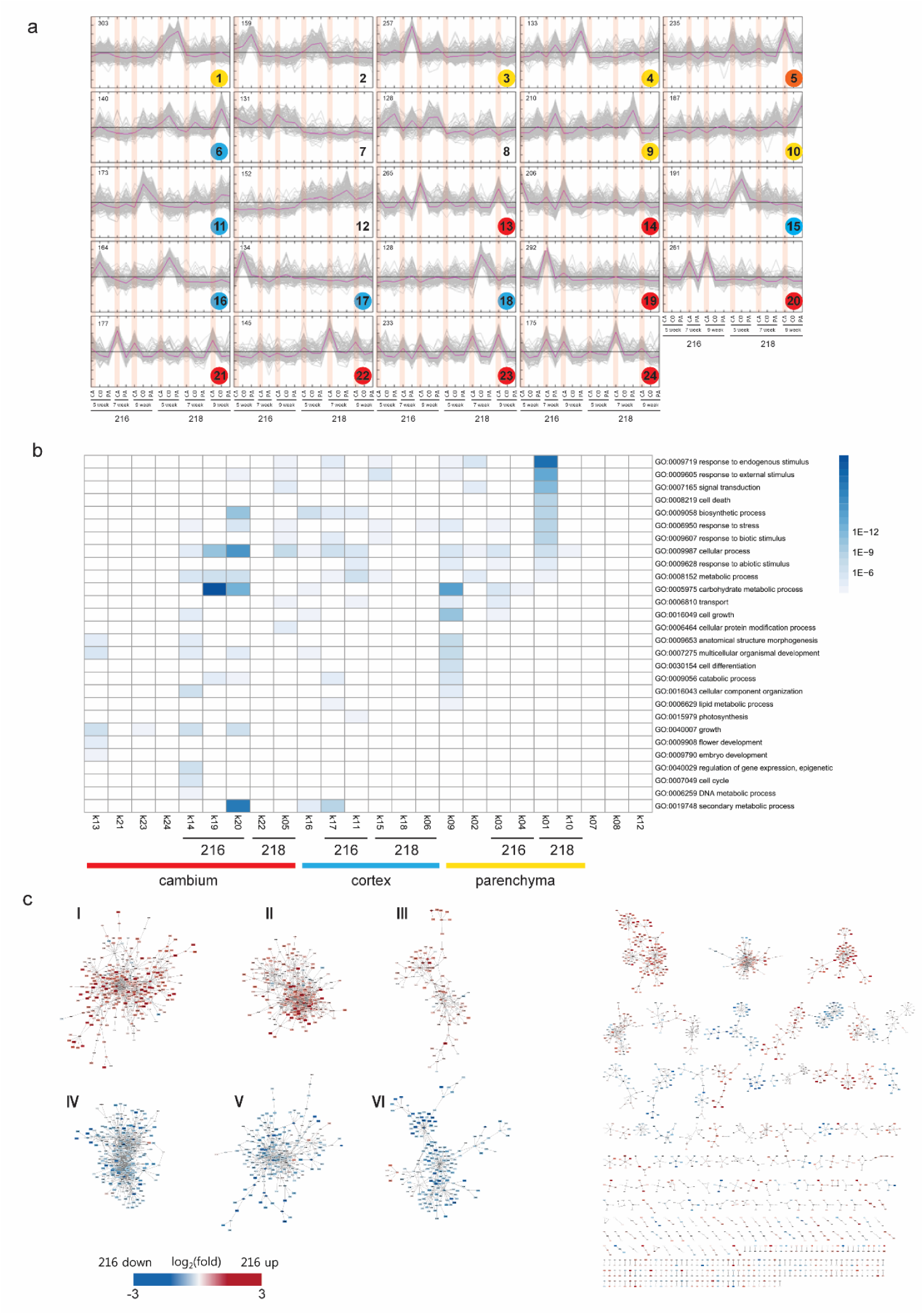
Cluster expression patterns of DEGs, GO Slim enriched biological processes and cross-species co-expression gene regulatory network (GRN) analysis of cambium-enriched TFs from Arabidopsis and radish. **a** k-means clustering of DEGs. 4,602 DEGs were grouped into 24 clusters, and color coded based on their tissue-preferential expression patterns. Vertical lines in red indicate cambium sample positions. **b** GO Slim categories of each co-expressed clusters were presented as a heatmap with color gradient based on the corrected p-value (p-value cutoff 0.001). Colors: red (cambium), blue (cortex), and yellow (parenchyma). **c** Summary of co-expression GRN analysis using Arabidopsis and radish data, in which six major GRN clusters were identified in radish. Log_2_(fold change) was used to construct the heatmaps.

Recently, we and colleagues reported comprehensive cell-type-specific expression data in the Arabidopsis roots that include the early stage of cambia (Zhang et al., 2019). Using this information, we further asked if relationships between the secondary growth and stress responses implicated from the radish data are evolutionarily constrained. We assembled cambium-enriched TFs either in Arabidopsis or in radish and used these TFs for constructing separate co-expression gene regulatory networks (GRNs) based on the radish root expression data and the Arabidopsis root expression data (Zhang et al., 2019). From two networks, shared edges between orthologous gene pairs were selected, and GRN was constructed. Based on the GRN, cambium-specific GRN was constructed, and then clusters were identified (Fig. 4c; Supplementary Fig. 14, Supplementary Tables 6A-C). Six major GRN clusters were identified in radish: GRNs I to III were composed of genes that were up-regulated in the fast-growing line 216, whereas GRNs IV to VI were with genes down-regulated. Top GO terms for the GRNs I to III were the xylem development, sucrose transport, and response to chitin in (Supplementary Fig. 15; Supplementary Table 6D). GRN V with suppression of expression in line 216 was significantly overrepresented by genes involved in multiple biotic/abiotic stress responses. These cross-species analyses further support that the GRN with genes involved in the stress responses might be an important contributor to the root secondary growth.

### Functional analyses of stress response network find ERF-1 as a key checkpoint for the cambium function

Cross-species analyses indicated the conservation of a GRN composed of stress response TFs and its association with the root secondary growth. To further characterize this aspect, we selected stress response TFs whose expression is highly enriched in the early cambia in Arabidopsis roots: *ERF-1*, *ERF2*, *MYB15*, *STZ*, *WRKY18*, *WRKY33, WRKY46*, and *ERF105* (Supplementary Table 7A). All of these were up-regulated in the cambia of line 218 (Supplementary Table 7B). In addition, most except for *MYB15* and *ERF105* were highly connected in conserved GRN cluster V (Supplementary Table 6C). Our analysis of root cross sections right below hypocotyls of 10 day-old seedlings for available knockout mutants, *erf-1*, *erf2*, *myb15*, *wrky33*, and *stz*, indicated that *erf2*, *myb15*, and *stz* have fewer cells derived from cambia than the wild type (Supplementary Fig. 16a).

Next, as the secondary growth regulators, we selected *WOX4*, *WOX14*, *PXY*, *SCL7*, *ASL9*, and *LHW*. The first five genes have been reported to regulate cambium activities by forming a complex GRN and show the mild defect in the root secondary growth when they were mutated (Supplementary Fig. 16b). We added *LHW* to further understand how it interacts with other characterized regulators and stress response factors.

To infer *in vivo* GRN, seeds of each selected loss-of-function (LOF) mutant were germinated and grown vertically on MS media for 10 days and then 1cm long root segments right below the hypocotyls were collected for quantitative RT-PCR experiments. We chose 10 day after seed planting for this experiment to capture the effect of gene perturbation at the time point when the cambium is just established. If the expression of a selected gene was up- or down-regulated by more than 1.5 fold, the target gene was considered to be downstream of the gene knocked-out (Supplementary Tables 8A and B). One network visualized with Cytoscape (Shannon et al., 2003) in Fig. 5a shows the edges in red when the edge supported by differential gene expression overlaps with the DAP-seq result (O’Malley et al., 2016). In this, we found that STZ differentially regulates *PXY*, *WOX4, WOX14*, and *ASL9* potentially via binding on their promoters while *STZ* and *ERF-1* control each other likely via direct regulation. *LHW* was also affected by *STZ* and *ERF-1* while it, in turn, regulated multiple stress response TFs. These results indicate that stress response TFs and the secondary growth regulators intimately regulate each other in the highly complicated GRN. From the two independent *in vivo* GRN analyses, we searched for genes consistently serving as key nodes in the networks by measuring betweenness and radiality and found *ERF-1* with highest scores of betweenness (Fig. 5b and Supplementary Fig. 17).

**Figure 5.**
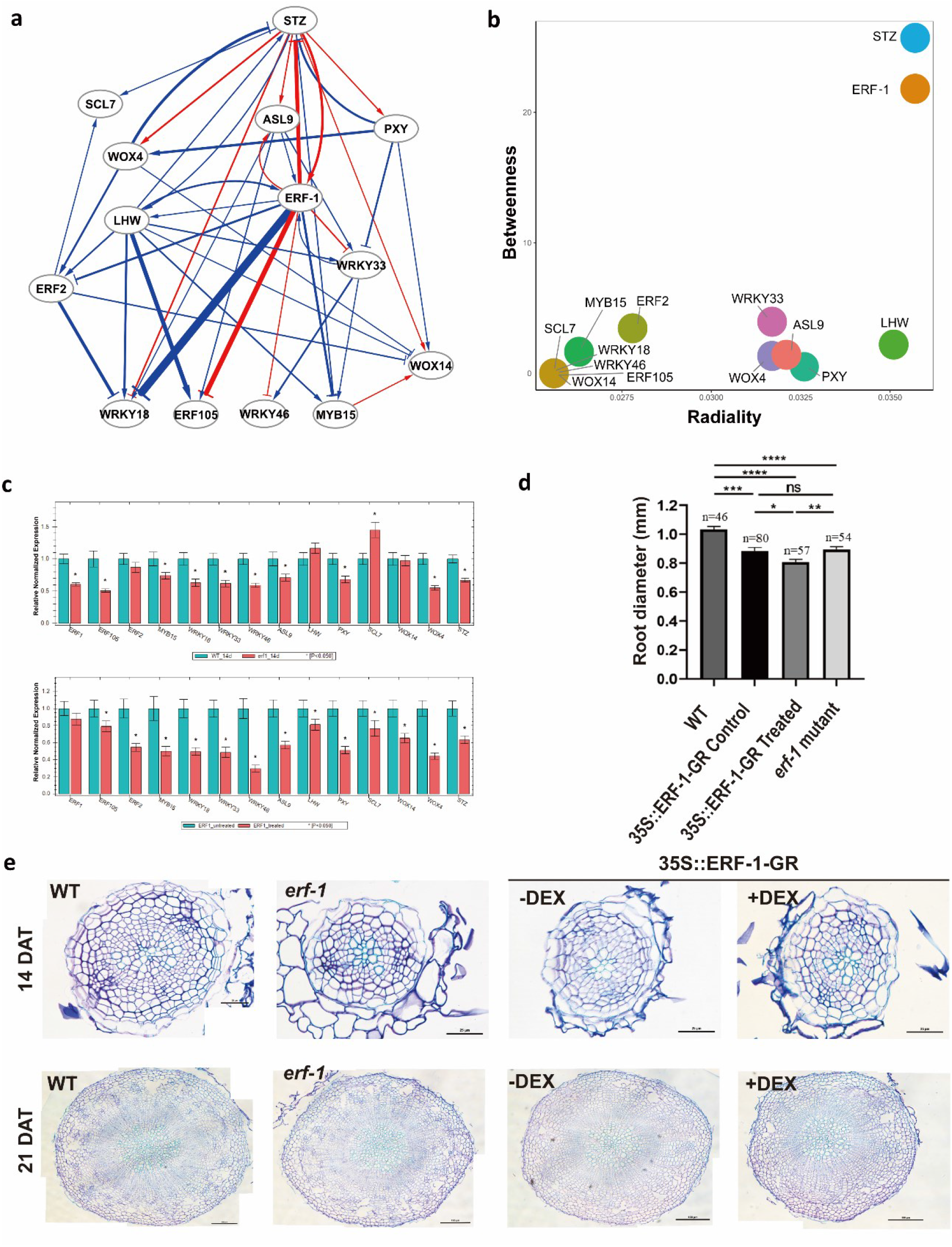
Functional analyses of stress response network. **a** A directed *in vivo* gene regulatory network constructed using the expression data obtained from perturbation experiments of 10d old knock-out mutant plants. Differentially expressed genes were defined at a |fold change| >1.5 and p <0.05). Arrow direction indicates source - target, while arrow thickness denotes the magnitude of gene regulation (fold change). The red edges indicate those edges that were supported by the DAP-Seq result (O’Malley et al., 2016). **b** Scatter plot of radiality vs. betweenness of key nodes in the network presented in a. **c** The expression level of selected genes in *erf-1* mutant (loss-of-function) and *35S∷ERF-1:GR* (gain-of-function) lines measured by qRT-PCR using 14d-old plants. Data were compared against WT (col-0) and untreated (*35S∷ERF-1:GR* without DEX treatment). Three technical replicates were run for each sample. Expression values were quoted as Mean ± SEM. Asterisks; statistical significance based on Student’s t-test (p < 0.05). **d-e** Comparison of root diameters and the xylem areas of 2- and 3- week old Arabidopsis plants with *ERF-1* loss-of-function or gain-of-function. WT (col-0) and *35S∷ERF-1:GR* without DEX treatment were used as controls, respectively. Data were quoted as Mean ± SEM. Asterisks; statistical significance based on ANOVA, ****p < 0.001, ***p < 0.01, **p < 0.05, *p < 0.1 and non-significant (ns). n denotes the number of samples.

In our GRN, *ERF-1* seemed directly connected with *STZ*, which was reported to enhance the tolerance of plants to abiotic stress when it is up- or down-regulated (Mittler et al., 2006). We thus asked how *ERF-1* behaves when it is up- or down-regulated. To find the effect of gain-of-function ERF-1, we used *35S∷ERF-1:GR* (glucocorticoid receptor) plants in which ERF-1:GR activity in the nuclei can be induced via binding of dexamethasone (dex) to the GR. When we compared the expression of selected genes in the presence of dex to the one without dex, we found the down-regulation of all the stress response TFs and the secondary growth regulators (Fig. 5c). This trend was similarly observed in the *erf-1* mutant, suggesting that balancing *ERF-1* level is important for the root secondary growth. Because both up- and down-regulating *ERF-1* result in the suppression of secondary growth regulators, we predicted that both cases would result in a decrease in root secondary growth. Indeed, both 2- and 3-week old Arabidopsis plants with *ERF-1* loss- or gain-of-function showed the decrease in the root diameters and the xylem areas in comparison to the control (Fig. 5d, e). These collectively indicate that ERF-1 might serve as a key node in the secondary growth GRN that integrates the growth/development program and the stress sensing program.

## Conclusion

In summary, our time-course transcriptome profiling of cambium and its neighboring tissues in two radish inbred lines provides rich information about transcriptional dynamics during the storage root growth. Comparative analysis based on the findings in Arabidopsis indicates evolutionary conservation of core regulatory processes in the cambium. In this, we found many cambium-enriched regulators whose functions in the secondary development have not been reported. How these are wired with regulators characterized in this and other studies is yet to be identified.

One major interest in the understanding of secondary development is to apply this to producing crops with enhanced yields or resistance to changing environments. Our comparison of transcriptome dynamics in inbred lines with contrasting growth and yields indicated that a GRN enriched with genes involved in sensing and responding to environmental changes intimately control the storage root growth. Considering that this GRN is conserved in the Arabidopsis root cambium as well, it might function as a key contributor to the secondary growth in general to fine-tune the growth in an ever-changing environment. The potential direct regulation of *PXY*, *WOX4*, and *WOX14* by STZ based on DAP-seq result further supports this idea. With a more detailed understanding of cambium GRNs, we might be able to design the crops with sustainable yields. In that context, our investigation provides a platform for future designing schemes.

## Methods

### Plant materials and growth

Radish inbred lines 216 and 218 were obtained from National Institute of Horticultural and Herbal Science (NIHHS) of Republic of Korea. Arabidopsis perturbation lines, *lhw* (SALK_079402), *pxy* (SALK_009542), and *wox4* (GABI462G01) were generously provided by Drs. Fukuda, Kondo, and Ohashi Ito at Tokyo University, and *erf-1* (SALK_036267), *wrky33* (SALK_006602), *stz* (SALK_054092), *erf2* (FLAG_314D04), *35S∷ERF-1:GR* lines (Van den Broeck et al., 2017) were by Dr. Inze at Ghent University. *myb15* (SALK_151976) was from ABRC and *asl9* mutant was developed in the lab using CRISPR-CAS9 system (Zhang et al., 2019).

Radishes were grown in a pot (20×20×20cm) at 22°C in long-day conditions (16 hour light/8 hour dark, 156 μmolm^−2^s^−1^) for 5, 7, and 9 weeks respectively before sample preparation. For the cambium analysis of Arabidopsis grown for more than 3 weeks, plants were grown in a tray with 32 pots under 16 hour light/8 hour dark and 125 μmolm^−2^s^−1^. Perturbation lines for analyses and Col-0 were planted randomly and grown in the same tray to avoid any bias caused by subtle differences in growth condition. For qRT-PCR analyses, Arabidopsis seeds were surface-sterilized and then plated on Murashige and Skoog (MS) medium supplemented with 1% Sucrose and 1% Agar. Seeds on MS plates were cold-treated at 4°C for 2 days, transferred to a growth chamber (22°C, 16 hour day and 8 hour night), and grown vertically.

### Embedding and sectioning for laser capture microdissection (LCM)

To prepare tissue sections, sample preparation protocol described by Thiel *et al* (Thiel et al., 2011) was used with minor modifications. Briefly, tissue block (1.2×1.2×1.2 cm^3^) containing cambium was obtained from the thickest part of a root. Dissected tissue block was immediately fixed in ice-chilled Famer’s fixative (75% ethanol, 25% glacial acetic acid), and stored overnight at 4°C. The fixed tissue samples were gradually dehydrated with ethanol and embedded with Steedman’s wax (9:1 (w/w) of polyethylene glycol 400 distearate and 1-hexadecanol). To obtain tissue sample slides for LCM, about 2mm of tissue sections were trimmed off (100 steps 20μm-thickness using a rotary microtome) to reach to the inner part of the tissue block, and then 20μm-thick tissue sections were prepared on an Arcturus PEN membrane glass slide.

### LCM and RNA extraction

Before LCM, Steedman’s wax was removed by incubating the slide in ethanol for 10 min twice, dried for 5 min, and immediately processed for LCM using an Arcturus Veritas LCM microdissection system. Areas of interest within the tissue were marked using the drawing tool in the Arcturus software to cut-out tissues using a UV laser (355nm), and the dissected tissues were collected onto the Arcturus Capsure Macro LCM cap. Total RNA was extracted with the Arcturus PicoPure RNA isolation kit following the manufacturer’s instructions. RNA quality was evaluated using RNA Pico chip on an Agilent 2100 Bioanalyzer system (NICEM, Seoul National University).

### Library construction and Illumina sequencing

RNA-Seq libraries were prepared using the Nugen Ovation RNA-Seq System 1-16 for Model organisms (Arabidopsis). An aliquot of 10 ng of each total RNA sample was used as a starting material, and final sequencing libraries were prepared by amplifying the constructed libraries using PCR with empirically determined amplification cycles. Paired-end sequencing data were obtained with Illumina HiSeq4000 (CLC Genomics and Epigenomics Core Facility, Weill Cornell Medical College).

### Read processing, transcriptome assembly and novel gene identification

Raw RNA-Seq reads were processed to trim the adapter and low-quality sequences using Trimmomatic (Bolger et al., 2014). After trimming, reads shorter than 80 bp were discarded. The resulting reads were aligned to ribosomal RNA (rRNA) database (Quast et al., 2013) using Bowtie (Langmead et al., 2009) allowing up to 3 mismatches. The aligned reads were discarded and the remaining high-quality cleaned reads were assembled *de novo* using Trinity (Grabherr et al., 2011) with “min_kmer_cov” set to 5. The resulting assembled contigs were then compared to the GenBank Nucleotide (nt) and non-redundant proteins (nr) databases using the BLAST+ program, and those having hits only to sequences from viruses, bacteria, and archaea were discarded. To remove redundancies in the Trinity-assembled contigs, the contigs were further processed using CD-HIT (Li and Godzik, 2006) with minimum sequence percent identify set to 95%.

To identify novel radish genes, the assembled contigs were mapped to the radish reference genome (http://radish-genome.org/) using GMAP (Wu and Watanabe, 2005) with “npaths” set to 0. Contigs overlapping with known gene regions in the reference genome were discarded. The coding potential of the remaining contigs was calculated using Coding Potential Calculator (CPC) (Kong et al., 2007), and contigs with CPC score > 0 and containing an open reading frame (ORF) > 300 bp were identified as novel radish genes. To annotate the newly identified genes, their protein sequences were compared to GenBank nr, the Arabidopsis protein and UniProt (Swiss-Prot and TrEMBL; http://www.uniprot.org/) databases using DIAMOND (Buchfink et al., 2015) with ‘-evalue’ set to 1e-4, as well as the InterPro database using InterProScan (v5.19-58.0) (Jones et al., 2014). Arabidopsis genes hit with the lowest e-value were considered the putative ortholog of radish transcripts. Those used for cross-species comparison are listed in the Supplementary document Table 5. GO annotations were obtained using Blast2GO (version 2.5.0) (Conesa and Gotz, 2008) based on the BLAST results against the GenBank nr database and results from the InterProScan analysis.

### LincRNA identification and analysis

To identify long intergenic noncoding RNAs (lincRNAs), a reference-guided transcriptome assembly was performed using Cufflinks (Trapnell et al., 2010) with high-quality cleaned RNA-Seq read pairs that were aligned to the radish reference genome. Expression of the assembled transcripts was measured and normalized to FPKM using the Cuffnorm program in the Cufflinks package. The coding potential of the assembled transcripts was calculated using CPC (Kong et al., 2007). LincRNAs were defined as transcripts of a minimum length of 200 nt, having CPC score of <-0.5 and located in the intergenic regions, with a distance >500 bp to the nearest radish protein-coding genes. These lincRNAs were also confirmed to be devoid of any ORFs >100 aa and to be expressed at an FPKM >0.1. Tissue specificity of lincRNAs was calculated for all tissues and developmental stages by employing the entropy-based measure (Cabili et al., 2011), which defines the expression patterns using Jensen-Shannon (JS) divergence score. The analysis and validation of lincRNAs are available in the supplementary document and Supplementary Table 9A.

### Differential gene expression analysis and clustering

The high-quality cleaned paired-end RNA-Seq reads were aligned to the radish genome *Rs* 1.0 (http://radish-genome.org/) and aforementioned novel genes using HISAT (Kim et al., 2015) allowing up to 3 edit distances. Following alignments, raw counts for each gene were derived and normalized into the number of fragments per kilobase of exon per million mapped fragments (FPKM). For differential gene expression analysis, lowly expressed genes were filtered out based on the total read counts across all samples less than 12. DESeq2 (Love et al., 2014) was used to obtain differentially expressed genes with a cut off of fold change >2 and False Discovery Rate (FDR) <0.05. Sample distance matrix and PCA analysis were performed using regularized log-transformed count data. For PCA analysis, the top five thousand most variably expressed genes were used. Pheatmap R package (https://cran.r-project.org/web/packages/pheatmap/) was used for data visualization. Cluster analysis was conducted using the MeV software (Saeed et al., 2003), and k-means clustering was performed with k=24. GO terms enriched in the set of differentially expressed genes and clustered genes were identified using GO∷TermFinder (Boyle et al., 2004).

### RNA *in situ* hybridization

Root tissues from radish inbred lines 216 and 218 grown for 5, 7, or 9 weeks in long-day conditions were fixed in cold 4% formaldehyde. Thin root tissue sections were prepared from Paraplast embedded tissue blocks for RNA *in situ* hybridization using DIG-labeled riboprobes. Detailed methods and protocols were provided in the supplementary document and Supplementary Tables 9B-E.

### Arabidopsis root imaging

For Arabidopsis root imaging and phenotyping, 4 week-old soil-grown Arabidopsis root segment was cut about 500 μm from the root-hypocotyl junction and fixed in 4% formaldehyde. Tissue was embedded in Technovit 8100 (Kulzer, Germany), sectioned at 6um, and imaged under Nikon ECLIPSE N*i* microscope.

### Arabidopsis and radish cross-species network analysis

Pearson’s correlation coefficients (PCCs) were calculated using root-specific expression data to find the relationships between TFs and target genes (TGs) in Arabidopsis and radish, respectively. To obtain strongly associated TF-TG pairs, PCC cut off was set to an absolute value of 0.67 (Ahn et al., 2017) and then a root-specific TF GRN for each species was constructed. To identify conserved functional relationships between two species in terms of secondary growth and stress response, cross-species GRN was assembled by intersecting the shared edges of two networks. To focus on the cambium-related genes in radish inbred lines, the property of expression patterns, such as consistently up-regulated or down-regulated at three growth stages, was characterized. Specifically, the cambium-specific template GRN (or template GRN) was constructed by profiling cambium time-course transcription data for each gene. On the template GRN, gene expression data of line 216 and 218 with each growth stage were mapped. Then, inbred-line difference GRNs were instantiated for capturing expression differences between line 216 and 218. Community detection analysis was performed (Su et al., 2010) and six major GRN clusters were identified by a statistical test with gene expression and GO terms. Detailed methods were provided in the supplementary document.

### Perturbation experiments and quantitative RT-PCR

LOF mutant lines were obtained for nine genes including *ASL9*, *WOX4*, *PXY*, *LHW*, *ERF-1*, *ERF2*, *MYB15, WRKY33* and *STZ* by CRISPR-CAS9 and T-DNA insertion. All T-DNA insertion lines were confirmed by PCR and Sanger sequencing. LOF lines were grown on MS medium for 10 days after transfer (DAT) from 4°C to the growth chamber, while *35S∷ERF-1:GR* line was grown MS medium for 9 DAT, and then subjected to 10μM of dex treatment for 5 days.

For qRT-PCR, primary root segments undergoing secondary development (~1 cm below hypocotyl) pooled from at least three plates (at least 30 plants) were used in RNA extraction employing Qiagen RNeasy plant mini kit. The first-strand cDNA was synthesized according to the protocol as for lincRNA validation section (see Supplementary Methods). cDNA template was diluted 25 times and subsequently used for qRT-PCR employing iQTM SYBER Green supermix (Bio-Rad) on a BioRad CFX96 Real-Time PCR machine, with the following conditions: 95°C for 3 min, followed by 48 cycles (95°C for 30 sec, 57°C for 10 sec, 72°C for 30 sec). Gene-specific primers were designed by NCBI Primer-BLAST (Ye et al., 2012). List of primer sequences is provided in Supplementary Table 9F. The expression level of each gene was normalized against that of the reference gene GAPDH, and compared against that of the control wild type sample (Col-0). Three technical replicates were run for each sample. Data were analyzed using Student’s *t*-tests Bio-Rad CFX Maestro software 1.0. Expression values were quoted as Mean ± SEM, and *p* <0.05 was considered statistically significant.

## Data availability

All RNA-Seq data including 33 paired-end read files generated for this study have been deposited publicly in NCBI SRA database under BioProject PRJNA475856, Study Accession Number SRP150507. All other relevant data that supports the finding in this study are included within the article and supplementary files.

## Acknowledgements

We thank Drs. Fukuda, Ohashi-Ito, and Kondo at University of Tokyo and Dr. Inze at Ghent University for providing mutant seeds. We are grateful to Dr. Suhyoung Park at National Institute of Horticultural and Herbal Science for providing two radish inbred lines 216 and 218. We also thank Lee lab members for assisting experiments at various stages.

This work is supported by National Research Foundation of Korea (NRF-2018R1A5A1023599) and Korea Institute of Planning and Evaluation for Technology in Food, Agriculture, and Forestry (IPET) through Golden Seed Project (213006-05-3-SBK30), funded by Ministry of Agriculture, Food and Rural Affairs (MAFRA), Ministry of Oceans and Fisheries (MOF), Rural Development Administration (RDA) and Korea Forest Services (KFS) to J-Y.L. G.C., A.C.A.F., J.H., and N.V.H. were supported by Brain Korea 21 Plus Program.

## Author contributions

G.C. contributed to radish tissue preparation, RNA-Seq library construction, DEG analysis, and network analysis; N.V.H. contributed to lincRNA analysis and RT-PCR validation, and *in vivo* GRN construction; Y.Z. contributed to transcriptome assembly, annotation, and gene ontology analysis; A.C.A.F. contributed to RNA *in situ* hybridization; J.H. contributed to Arabidopsis mutant analysis; I.S., H.A., S.K. contributed to cross-species comparison of gene regulatory networks; J-Y.L. contributed to managing and designing experiments and analyses; G.C., N.V.H.,Y.Z., A.C.A.F., Z.F., S.K. and J-Y.L. contributed to manuscript writing. All authors read and approved the final manuscript.

## Correspondence

Correspondence to Ji-Young Lee

## Supplementary data

**<u>Supplementary document and Figure file</u> contains the following information:**

**Supplementary materials and methods**

**Supplementary figures 1-17**

- **Supplementary Figure 1.** Radish samples for RNA-Seq experiment and RNA-Seq library construction workflow.
- **Supplementary Figure 2.** *De novo* transcriptome assembly and transcript annotation workflow.
- **Supplementary Figure 3.** Cambium DEGs shared in all growth stages and inbred lines.
- **Supplementary Figure 4.** RT-PCR validation of 32 selected putative lincRNAs identified in this study.
- **Supplementary Figure 5.** Heatmaps of expression of lncRNAs vs neighboring coding transcripts.
- **Supplementary Figure 6.** Heatmaps of expression of natural antisense transcripts (NATs) vs *trans*-coding transcripts.
- **Supplementary Figure 7.** Secondary structure of 32 radish miRNA precursors identified in this study.
- **Supplementary Figure 8.** Heatmaps of expression of miRNA-encoding lincRNAs and target genes.
- **Supplementary Figure 9.** Row normalized expression of genes selected for RNA *in situ* hybridization.
- **Supplementary Figure 10.** RNA *in situ* hybridization with sense probes.
- **Supplementary Figure 11.** Auxin pathways.
- **Supplementary Figure 12.** Cytokinin biosynthesis and signaling pathway.
- **Supplementary Figure 13.** BR and GA pathway.
- **Supplementary Figure 14.** Workflow for Arabidopsis and radish cross-species network analysis.
- **Supplementary Figure 15.** Conserved radish GRN clusters showing the relative expression of each node in the cambium in comparison between 216 and 218.
- **Supplementary Figure 16.** Root cross-section images for knockout mutant lines used in the *in vivo* GRN analyses.
- **Supplementary Figure 17.** GRN network and scatter plot of radiality vs. betweenness constructed from qRT-PCR data from the second batch of perturbation experiment

**<u>Supplementary Table 1</u> contains the following tables:**

- **Table S1A.** Summary statistics of sequencing read data used in this study
- **Table S1B.** List of novel genes identified in this study
- **Table S1C.** Raw count tables for all transcripts from 33 samples
- **Table S1D.** Mean FPKM for all transcripts from 17 samples

**<u>Supplementary Table 2</u> contains the following tables:**

- **Table S2A.** List of DEGs identified for cambial tissue samples
- **Table S2B.** Total mean FPKM for DEGs identified in cambial tissue samples

**<u>Supplementary Table 3</u> contains the following statistic tables for lincRNA identification analysis**

- **Table S3A.** Summary of pre-processing and read mapping for lincRNA analysis
- **Table S3B.** Sample correlation table
- **Table S3C.** FPKM values for lincRNAs

**<u>Supplementary Table 4</u> contains the following tables:**

- **Table S4A.** Pearson’s correlation of lincRNAs and their nearby protein-coding transcripts
- **Table S4B.** Pearson’s correlation of identified NATs and their target protein-coding transcripts
- **Table S4C.** Identified miRNA encoding lincRNAs in this study
- **Table S4D.** Pearson’s correlation of identified miRNA-encoding lincRNAs and their target protein-coding transcripts

**<u>Supplementary Table 5</u> contains 24 GOSlim tables from k01-k24 and their statistics**

**<u>Supplementary Table 6</u> contains the following tables:**

- **Table S6A.** Node information for conserved GRN clusters
- **Table S6B.** Edge information for GRN clusters
- **Table S6C.** Transcription factors in GRN clusters
- **Table S6D.** GO enrichment analysis of GRN clusters

**<u>Supplementary Table 7</u> contains the following tables:**

- **Table S7A.** Expression of selected stress response TFs and secondary growth regulators which were highly enriched in the early cambia in Arabidopsis roots
- **Table S7B.** Expression of conserved stress response TFs and secondary growth regulators in radish roots

**<u>Supplementary Table 8</u> contains the following tables:**

- **Table S8A.** Expression of selected stress response TFs and secondary growth regulators in 10-day knockout mutant lines by quantitative RT-PCR (batch 1)
- **Table S8B.** Expression of selected stress response TFs and secondary growth regulators in 10-day knockout mutant lines by quantitative RT-PCR (batch 2)

**<u>Supplementary Table 9</u> contains the following tables:**

- **Table S9A.** Primers used for validations of 40 selected lincRNAs by RT-PCR
- **Table S9B.** Arabidopsis ortholog IDs of radish genes selected for *in situ* hybridization
- **Table S9C.** Primers used for cloning
- **Table S9D.** Primers used for probes
- **Table S9E.** Radish gene symbols and arabidopsis orthologs
- **Table S9F.** Primers used in quantitative RT-PCR

