## Supplementary Document and Figures for "Identification of conserved gene regulatory networks that integrate environmental sensing and growth in the root cambium"

- **Supplementary materials and methods**
- **Supplementary figures 1-17**
- **Supplementary document references**

**SUPPLEMENTARY DOCUMENT**

**SUPPLEMENTARY MATERIALS AND METHODS**

**Radish gene cloning and RNA *in situ* hybridization to validate RNA seq data**

**cDNA library preparation**

For preparing the cDNA library, 7-week-old radish root was grinded using a grinder and liquid nitrogen. Total RNA library was obtained by using the QUIAGEN RNeasy® Plant Mini kit obtaining a final volume of 50µl. For finally obtaining the cDNA library, a retro-transcription reaction was carried out. 2ng of the total RNA was mixed with 1µl of Oligo(dt)$_{16}$(10pM/µl), dNTP (10mM) 0,5µl reaching a total volume of 13µl. The mixture was incubated at 65˚C for % minutes and then 4µl of 5x 1$^{st}$ strand buffer was added, together with DTT 1µl, water 1µl and SuperScriptIII reverse transcriptase to a final volume of 7µl. The reaction was the incubated at 50˚C for an hour and 70˚C for 15 minutes fiving as a result 20µl of total cDNA library.

**Gene cloning**

A PCR reaction was done for this experiment with Phusion® High- Fidelity DNA Polymerase (NEW ENGLAND Biolabs). The PCR machine was set to work for 34 cycles, starting with a 30 seconds denaturalization at 98˚C, 30 second annealing temperature 3 degrees above the primer Tm, and an elongation at 72˚C which during 30 seconds for every kb of product.

The product was run through an electrophoresis gel 1% agarose to verify the absence of secondary products and purified by the Biofact kit HiGene Gel&PCR Purification System. The purified product was the inserted into pENTER/D-TOPO vector by TOPO$^{TM}$ Cloning reaction. For this reaction, 2µl of purified products together with 0.5µl of salt solution and 0.5µl of TOPO vector, was used. The mixture was incubated for 5 minutes at room temperature and transformed *into E. coli* by a 45 second 42˚C heat shock followed by a 2 minutes ice incubation. Colonies were selected using Luria-Bertani (LB) supplemented with the antibiotic kanamycin (50µg/ml).

**RNA *in situ* hybridization**

1. **RNA probe making**

PCR product was purified and concentrated into 100ng/µl. The transcription was carried out using T7 RNA polymerase reaction of the DIG labeling Kit from Sigma by adding 2n+n of acetylated BSA, 2n+n of transcription buffer, 2n+n of 10% Triton X-100, 2n+n of 10XNTP labeling mixture, with a final volume of 9µl. To the mixture described above, 9µl of purified PCR product and 2µl of T7 RNA polymerase was added.

The reaction was incubated for 37˚C for 3 hours and then DNA was removed from the mixture by adding 5µl of DNase buffer, 22µl of RNase Free water and 2µl of RNase-free DNase. The mixture was incubated 15 minutes at 37˚C. Finally, the probe was let to precipitate overnight at -80˚C using LiCl 6µl, 1µl of 0.5M EDTA and 180µl of absolute ethanol and resuspended in water.

**2. Tissue preparation**

The radish root was cut in the area were the secondary growth was maximum and it was immediately submerged in 4% paraformaldehyde fixative. The tissue was infiltrated using vacuum for 5 minutes 3 times and incubated overnight at 4˚C in fresh fixative.

The fixative was then removed and rinsed with Phosphate-Buffered Saline (PBS) 1X for 15 minutes four times. Tissue was then dehydrated using the following ethanol series for 1 hour: 25%, 50%, 75%, 100%, 100%, 100%, 75% ethanol/25% Histoclear®, 50% ethanol/50% Histoclear®, 25%ethanol/75% Histoclear®, 100%histoclear. 100%histoclear, 100%histoclear. Finally, half of the rube was filled with Paraplast® chips and left overnight at 58˚C for infiltration. For the following 4 days the Paraplast® was replaced twice a day and left at 58˚C. On the fifth day, the radish slices were oriented in a mold with molten Paraplast® left at room temperature to solidify. Finally, the tissue blocks were cut using a RM 2255 microtome (Leica) creating 15µm slices that were mounted with water on a TRUBOND® slide and left to dry overnight.

**3. Slide Pretreatment**

Slides were placed two times in Histoclear® for 10 minutes, then passed through ethanol series (100%, 100%, 95%, 85%, 70%, 50%, 30% in NaCl 0.85%) for 1 minute, NaCl 0.85% for 2 minutes, in 0.2M of HCl for 20 minutes, in water for 5 minutes, in 1XPBS for 2 minutes and treated with Pronase (0l.135mg/ml) for 28 minutes at 37˚C. Reaction was stopped by submerging the slides in Glycine 0.2% in PBS, followed by a 2 minutes submersion in 1X PBS and a 10-minute submersion in 4% paraforlmaldehyde fixative. Once, the fixation was finished, slides were rinsed for 2 minutes in 1x PBS, then submerged in NaCl 0.85% for 2 minutes and finally al the ethanol series were repeated backwards for dehydration of the tissue. Slides were left to dry in RNase free conditions for 1 hour.

**4. Pre-hybridization and hybridization**

For these steps, a chamber was prepared putting papers damp in 50% formamide covered in parafilm in a closed Tupperware to create a formamide atmosphere. Slides were placed in the chamber with 250µl of prehybridization solution (50% formamide, 1X salts, 1X Denhardt's, 200µg/ml of tRNA, 10U/ml of RNase inhibitor) covered by a microscope cover glassed. Slides were incubated 1 hour at room temperature ad 1 hour at 45˚C. During the prehybridization incubation, probes were incubated 1 minute at 80˚C to prevent secondary structures. Once the prehybridization incubation was finished, we added 250µl the hybridization solution (50% formamide, 1.25X salts, 12.55 dextran sulfate, 250µg/ml tRNA, 1.25X Denhardt's, 12.5 U/ml od RNase inhibitor, DEPC water) to the slides that contained 0.4µg/ml/kb of probe, we covered the slides with microscope cover slides and incubated the slides in the chamber for 24 hours at 45˚C.

**5. Post-hybridization washes**

After finishing the hybridization, the coverslips were removed by dipping the slides in pre-warmed 0.2X Saline Sodium Citrate (SSC) solution. Slides were the placed in a rack which stands in a jar with 0.2X SSC for 1 hour at 55˚C. Solution was later replaced and incubated for another hour. After finishing the washes, slides were rinsed in NTE solution (10mM of Tris pH 8.0, 5mM EDTA) and incubated for 30 minutes at 37˚C in a solution of 10µg/ml RNaseA in 0.5M NaCl, NTE. Slides were rinsed for 5 minutes in NTE for stopping the reaction and incubated one more hour in 0.2X SSC at 55˚C. Finally slides were rinsed in 1X PBS.

**6. Detection**

Sliced were placed in Blocking solution (100 mM Tris, 100mM NaCl, 1% blocking reagent od DIG Nucleic Acid detection kit, Roche) with gentle agitation for 45 minutes. Then, the blocking buffer was replaced by buffer A (100mM Tris, 100mM NaCl, 1% BSA, 0.3% Triton X-100) and slides were incubated for another 45 minutes. Once, the slides were ready, antibody conjugate from the DIG Nucleic Acid detection kit was spread on the slides in a 1:1000 ration in buffer A. Slides were incubated for 1 hour at room temperature in a chamber with high humidity. Finished this step, slides were washed in Buffer A with gentle agitation for 20 minutes 3 times and in detection buffer (100mM Tris pH 9.5, 100mM NaCl, 50mM $MgCl_2$) 2 times for 5 minutes. Finally, 500µl of color substrate (200µl of NBT/BCIP solution DIG Nucleic Acid detection kit in 10 ml of detection buffer and 100µl of levamisole) of the was added to the slides and covered with a coverslip and incubated at room temperature for 36 hours protected from the light.

**7. Imaging**

Images were taken in a Nikon e*clipse* N*i* light microscope. Morphology images were obtained by staining the sections with 0.05% toluidine blue (pH 4.4).

### LincRNA analysis and validation

#### LincRNA analysis

The potential roles of lincRNAs in regulating protein-coding mRNA were investigated as follows. First, the relationship of expression between lincRNAs and their nearest genes was assessed by Pearson's correlation analysis (Pearson, 1895). Second, we explored the possible role of lincRNAs as *trans* natural antisense transcripts (*trans*-NATs). To identify *trans*-NATs, we adopted the NATpipe (Yu et al., 2016) and queried the identified lincRNAs against a total of 50,027 protein coding sequences available for radish. These target sequences included 46,512 coding sequences obtained from the Radish genome Database (Rs 1.0 CDS, http://radish-genome.org) and 3,515 novel coding transcripts identified in this study. An all-v-all BLAST search (NCBI-BLAST 2.2.31) was performed to find matching NAT-CDS pairs, which then were parsed by a perl script *blastparser.pl* of the NATpipe. Putative *trans*-NATs were those derived from pairs which had complementary region longer than 50% of either sequence in each pair or consecutive complementary region ≥100 nt. For more than one trans-NAT matching the same target coding gene, the sum of FPKM values was calculated and used for Pearson's correlation analysis between the pairs.

Lastly, we searched for lincRNAs generating miRNAs and their potential target mRNAs. To this end, we adopted the miRNA prediction pipeline in CLC Genomics Workbench version 11 (CLC-GWB, CLC Bio-Qiagen, Aarhus, Denmark) employing a set of 4,933 non-redundant mature miRNAs from 48 selected plant species in mirBase release 22 (Kozomara and Griffiths-Jones, 2014) as reference. Precursor miRNA sequences were extracted from candidate lincRNAs based on two matching regions (maximum two mismatches) against the reference mature miRNAs. For lincRNAs showing only one matching region, we extracted sequences with 200 nt upstream and downstream to the mature miRNAs to include the second potential matching regions which could be more than two mismatches to the reference mature miRNAs. All extracted precursor miRNAs were subjected to secondary structure prediction using the RNAstructure web server (Reuter and Mathews, 2010) with default settings. We retained only sequences that folded into stem-loop hairpin structures based on their minimum free energy and base pairing probabilities. The expression of miRNAs was confirmed by read mapping of a set of radish small RNA sequencing data obtained from NCBI (accession number: SRR2729875). These steps were conducted to ensure that the identified miRNAs meet the recently updated criteria for plant miRNA annotation (Axtell and Meyers, 2018). Sequences from the 5' arm (normally corresponding to mature miRNA) and from the 3' arm (corresponding to passenger strand, miRNA*) of the miRNA duplex on the precursor miRNA were extracted for a target gene analysis using the plant small RNA target analysis server - psRNATarget ver2017 (Dai and Zhao, 2011) against the total 50,027 radish coding sequences (mentioned earlier). This assumed that some miRNA* could potentially function as mature miRNA, as observed in many studies (Guo and Lu, 2010; Okamura et al., 2008), and that the expression level of 5' arm sequence was shown not always higher than that of the 3' arm sequence by small RNA sequencing read mapping. A maximum expectation of 3.0, a size for complementarity scoring (HSP size) of 20, target accessibility-allowed maximum energy to unpair the target site (UPE): 25.0, flanking length around target site of 17 nt upstream and 13 nt downstream, a translation inhibition range of 10–11 nt and other default settings were used in target gene search. Relationship between miRNAs and their target mRNAs were assessed by Pearson's correlation analysis using the same approach as described above for NAT-target gene pairs.

**Validation of putative lincRNAs by RT-PCR**

To validate the putative lincRNAs identified in this study, we carried out RT-PCR for 40 selected lincRNAs including those ubiquitously expressed in two radish lines (216 and 218), those highly line-specific (expressed highly in one line in relation to the other line) and NATs that showed evolutionary conservation by comparing against *Arabidopsis* TAIR10 intergenic sequences ver20101028 (Swarbreck et al., 2008) using NCBI BLAST (E-value ≤1e-05). Total RNA samples were isolated from 7-week-old radish roots and treated with DNase I to remove any potential DNA contamination. The first strand cDNA was synthesized by priming with oligo-d(T) and transcribing from RNA with SuperScript™ III Reverse Transcriptase (Invitrogen, Carlsbad, CA, USA). The PCR conditions were as follows: 95°C for 3 min, followed by 40 cycles (95°C for 30 sec, 55°C for 40 sec, 72°C for 30 sec) and a final extension 72°C for 5 min. The radish *β*-ACTIN gene (Xu et al., 2012) were used as an experimental control, while RNA sample without reverse transcription step was used as negative control. Primers were designed by Primer3 (v.0.4.0) (Untergasser et al., 2012) using Tm (58-63°C), primer length (18-27 nt) and product size range (150-400 nt). All information related to selected lincRNAs and PCR primers is provided in Supplementary Table 4.

**Arabidopsis and radish cross-species network analysis**

To investigate relationship between the secondary growth and stress responses in radish, we wanted to utilize the known relationship between the secondary growth and stress responses in Arabidopsis. To achieve this goal, we performed cross-species network analysis by constructing and matching Arabidopsis and radish networks using both Arabidopsis and radish gene expression data. This network analysis was based on a two-step bioinformatics strategy of our previous work (Ahn et al., 2017) and the workflow was illustrated in Supplementary Figure 14.

- **Step 1**: Constructing a cambium-specific template gene regulatory network (GRN) using gene expression data of Arabidopsis and radish

- **Step 2**: Instantiating inbred-line difference GRNs and identifying differentially expressed GRN clusters

**Network analysis step 1: Constructing cambium-specific template GRNs from gene expression data of Arabidopsis and radish**

This step was to construct two cambium-specific TF-centric GRNs, one for Arabidopsis and another for radish, using root-specific gene expression data. For Arabidopsis GRN construction, Arabidopsis TF IDs were those that were curated in PlantTFDB (Jin et al., 2017) and the Arabidopsis root-specific microarray gene expression data were obtained from the literature (Zhang et al., 2019) (21 root-tissue locations, with one to four replicates, then a total of 66 microarray samples), and all pairwise Pearson's correlation coefficients (PCCs) were computed between TF-TF and TF-target gene (TG) on the Arabidopsis root-specific microarray gene expression data. Then, TF-TF or TF-TG pairs with high significance ($|PCC| > 0.67$) were selected to construct an "Arabidopsis root-specific GRN", where the threshold value of 0.67 was selected following the guideline in a previous work (Ahn et al., 2017).

A radish root-specific GRN was constructed in the same way as Arabidopsis network construction. Nucleotide sequences of all radish transcripts were compared to those of Arabidopsis transcripts, then each radish transcript was assigned an ID of Arabidopsis transcript that was the best-hit of the radish transcript among other Arabidopsis transcripts. All pairwise PCCs between TF-TF and TF-TG were computed on radish root-specific RNA-Seq data (two phenotypes, three tissues, three time-points with one or two replicates, then a total of 33 RNA-Seq samples). Then, TF-TF or TF-TG pairs with high significance ($|PCC| > 0.67$) were selected to construct a "radish root-specific GRN".

A cross-species network was constructed by integrating the two Arabidopsis and radish networks. Let $i_r$ and $j_r$ be two radish genes, and $i_A$ and $j_A$ be their best-hit Arabidopsis genes. The integrated network, called "cross-species GRN", consisted of the connections of $i_r$ and $j_r$ if $i_r$ and $j_r$ were connected in the radish network and $i_A$ and $j_A$ in the Arabidopsis network. The shared edges were selected if there were high co-expression associations between two genes on both Arabidopsis and radish root-spatial condition.

To determine shared edges more accurately, we need to consider fold change in addition to PCC since PCC considered the gradient of change only, not the absolute difference. For instance, two non-DEGs (differentially expressed genes) could provide a high PCC value. To overcome this issue, we conducted an additional filtering process based on the differential expression profiles for time-course gene expression data of cambium tissue type. We computed the fold change of expression of a gene, $e^{216}_i / e^{218}_i$, where the numbers, 216 and 218, were fast and slow growing inbred-line samples, respectively, and $i$ is the index of radish genes.

Then, we determined a differential expression profile based on the following criteria: 1 (up-regulated) if $e^{216}_i / e^{218}_i > 1.25$, -1 (down-regulated) if $e^{216}_i / e^{218}_i < 1/1.25$, 0 (no-changed) otherwise, where the threshold value of 1.25 was from the literature (Raices et al., 2017; Yoder et al., 2017). For example, if the differential expression profiles of gene *i* of three time points, 5, 7, and 9 weeks was [1, 0, -1], it meant the expression level of gene *i* on a 216 in-bred line sample was up-regulated, no-changed, down-regulated than that of 218 sample at 5, 7, and 9-week time-points on the cambium tissue. Then, we maintained edges of the cross-species GRN if two genes of an edge had similar differential expression profiles, such as [1, 1, 1] and [0, 1, 1]. Otherwise, edges were discarded if different profiles such as [1, 1, 1] and [0, -1, -1]. All remaining edges were used to construct the final network, called a "cambium-specific template GRN (or template GRN)".

**Network analysis step 2: Instantiating inbred line difference GRNs and identifying differentially expressed GRN clusters**

This step was to identify and characterize differentially expressed GRN clusters of the template network. First, sample-specific networks, called "inbred-line differential GRNs", were instantiated from the template network by mapping log gene expression differences between 216 and 218 inbred-lines, i.e., $log_2(e^{216}_i / e^{218}_i)$, to node values of the template network for each time point and each tissue type. Supplementary Figure 14 shows inbred-line differential GRNs with 17 TFs for cambium tissue with 5w, 7w, 9w, cortex tissue with 5w, 9w, and parenchyma tissue with 5w, 7w, 9w. Second, the GLay community detection algorithm (Su et al., 2010) partitioned the inbred-line differential GRNs into several gene clusters. Third, the six gene clusters that had large member genes (nGene ≥ 100) and were differentially expressed ($p <$ E-20, one-sample t-test with respect to zero mean) were selected as "major GRN cluster." Lastly, we characterized the six major GRN clusters by performing GO enrichment test for each GRN cluster using the David website (Huang et al., 2009) for each GRN cluster.

### SUPPLEMENTARY FIGURES

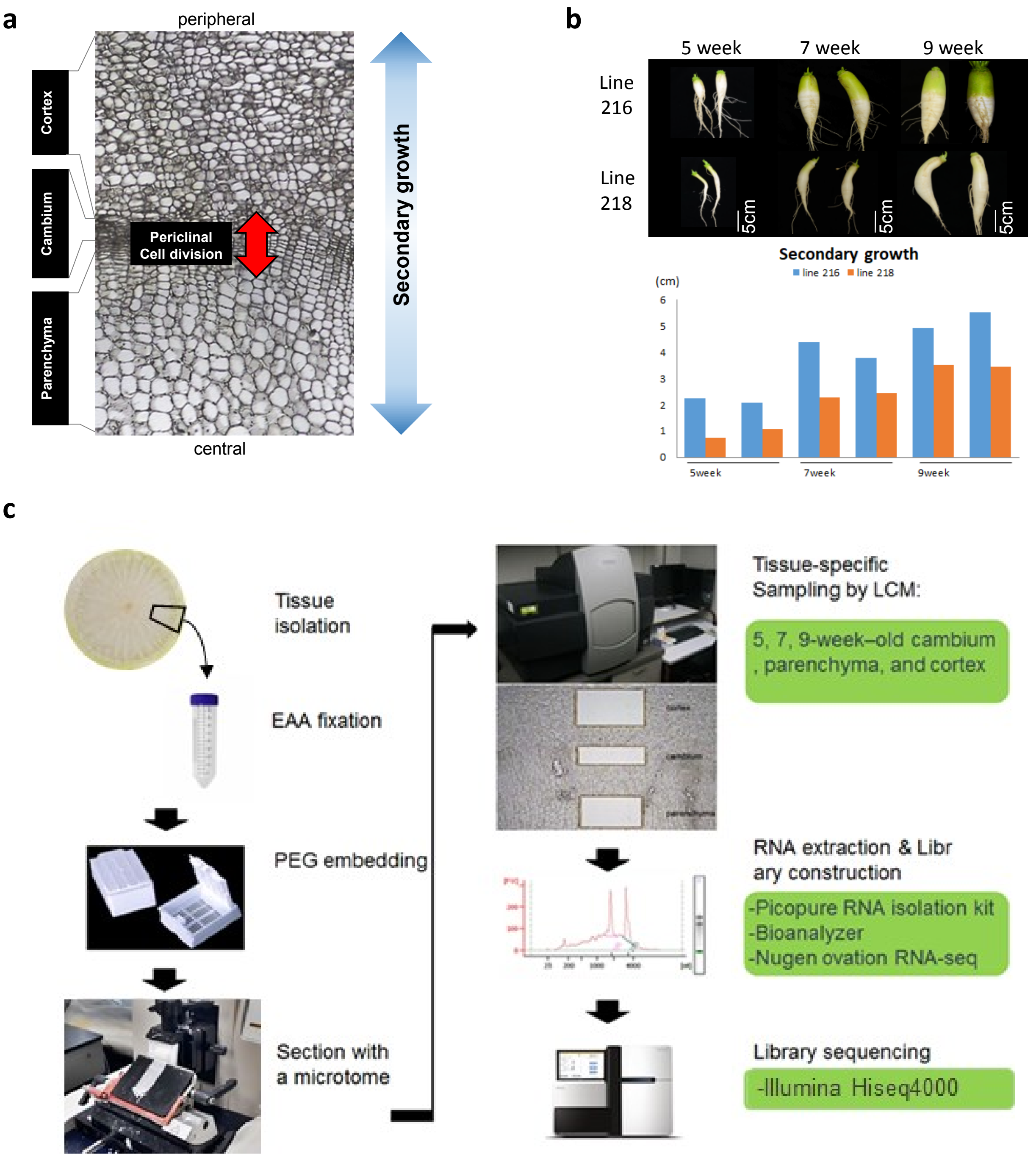


**Supplementary Figure 1. Radish samples for RNA-Seq experiment and RNA-Seq library construction workflow. a** Representative image showing radish root cross section used for RNA-seq library preparation. Bi-directional arrow in red shows cell division orientation in cambial cells during the secondary root growth. **b** radish inbred lines used for RNA-seq library preparation. Bar graph shows the root diameter of each radish root in the thickest region along the longitudinal axis. **c** RNA-seq library preparation workflow.

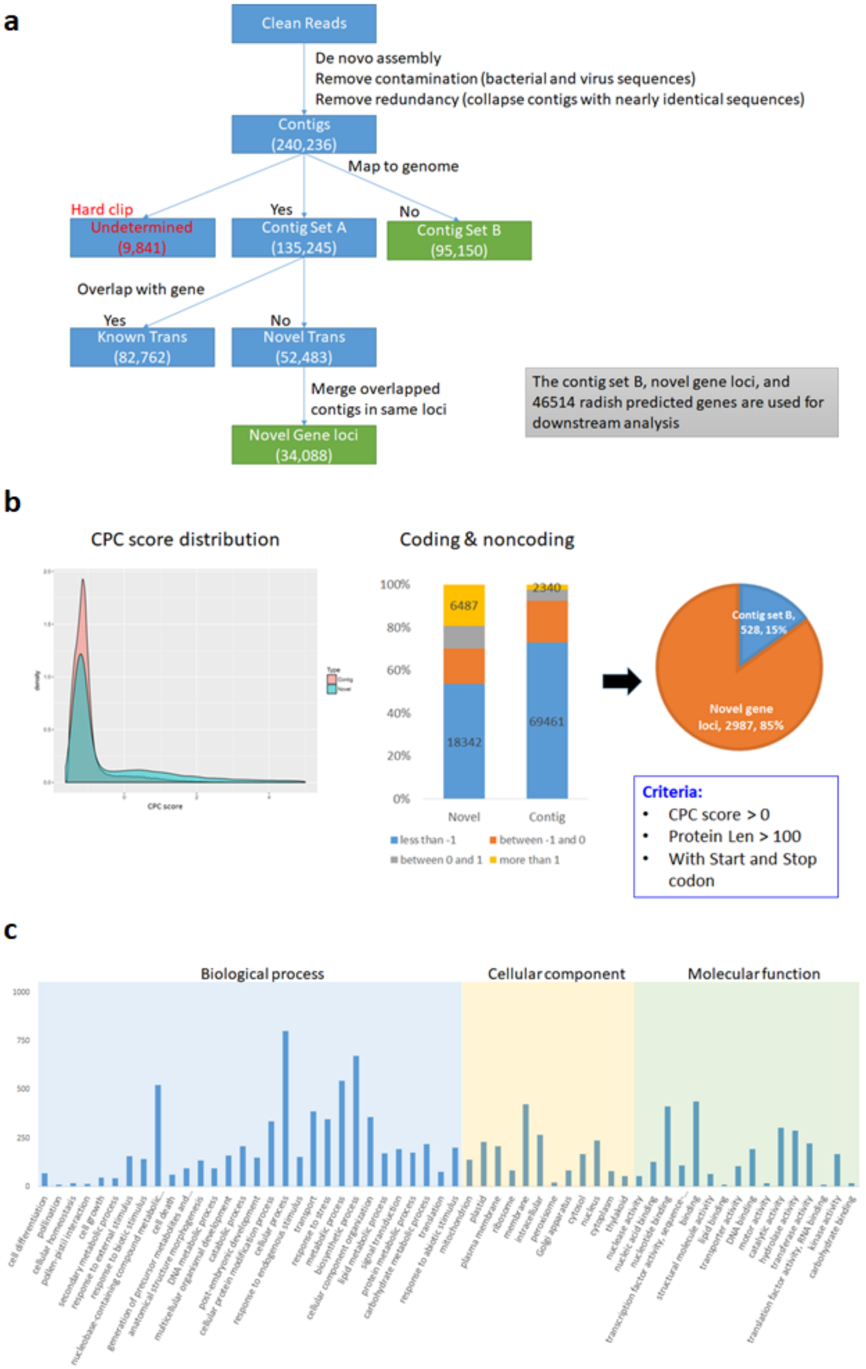


**Supplementary Figure 2.** ***De novo* transcriptome assembly and transcript annotation workflow.** a De novo assembly and transcript annotation pipeline. **b** Novel protein coding transcript assigned based on CPC score: density plot (left), bar plot (middle); pie chart (right) shows the percentages of novel transcripts. **c** GO categories of the novel transcripts

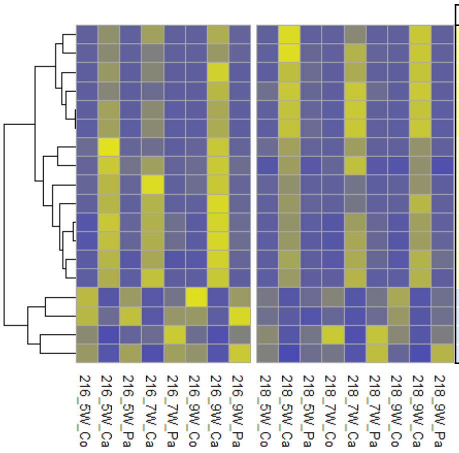


| Gene ID | ATH_ID | description |
|---|---|---|
| Rs160790.1 | AT4G20270.1 | BAM3, BARELY ANY MERISTEM 3 |
| Rs356140.1 | AT2G32860.2 | Beta glucosidase 33 |
| Rs219260.1 | AT3G01670.2 | SIEVE ELEMENT OCCLUSION-RELATED 2, SEOR2 |
| Rs041240.1 | AT4G37400.1 | Cytochrome P450 family 81 subfamily F polypeptide 3 |
| Rs475190.1 | AT1G63310.1 | Hypothetical protein |
| Rsnovo000323 | AT1G63310.1 | Hypothetical protein |
| Rs247050.1 | AT1G28290.1 | Arabinogalactan protein 31 |
| Rs286670.1 | AT3G09070.1 | OCTOPUS, OPS |
| Rsnovo000395 | AT2G24940.1 | Membrane-associated progesterone binding protein 2 |
| Rs454560.1 | AT1G71380.1 | Cellulase 3 |
| Rs159610.1 | AT4G22910.1 | FIZZY-related 2, CELL CYCLE SWITCH PROTEIN 52 A1 |
| Rs159660.1 | AT4G22910.1 | FIZZY-related 2, CELL CYCLE SWITCH PROTEIN 52 A1 |
| Rs022420.1 | AT1G28400.1 | GATA zinc finger protein |
| Rs045550.1 | AT4G32880.1 | Homeobox-leucine zipper protein ATHB-8 |
| Rs206730.1 | AT2G41560.1 | Autoinhibited Ca(2+)-ATPase isoform 4 |
| Rs403680.1 | AT5G22500.1 | Fatty acid reductase 1 |
| Rs249460.1 | AT1G09970.2 | Leucine-rich receptor-like protein kinase family protein, RLK7 |
| Rs304300.1 | AT3G23050.1 | Indole-3-acetic acid 7 |

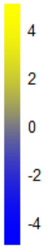


**Supplementary Figure 3. Cambium DEGs shared in all growth stages and inbred lines.** Row normalized expressed values were used for the heatmap visualization with Arabidopsis homologs and annotation information for the radish genes.

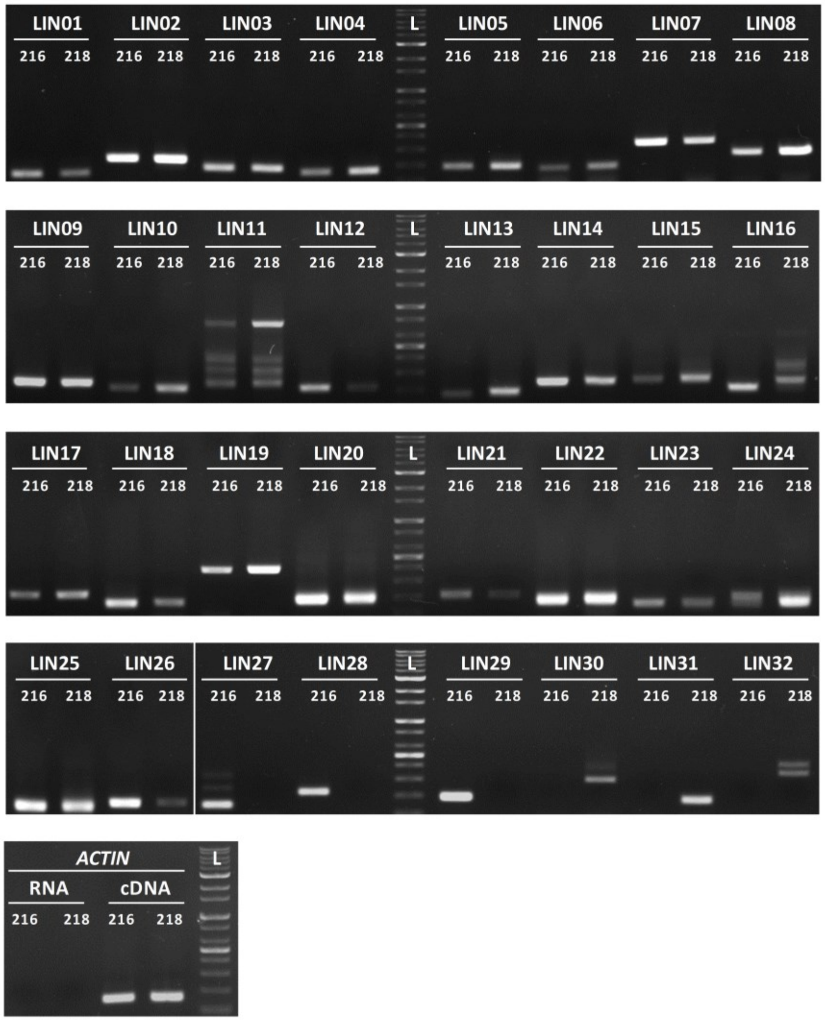


**Supplementary Figure 4.** RT-PCR validation of 32 selected putative lincRNAs identified in this study. Primer pair was designed for each of lincRNAs and run on RNA samples collected from 7 week-old plants of two radish lines, 216 and 218. Radish $\beta$-ACTIN gene was used as experimental control and DNase-treated RNA samples was used as negative control. All of 32 lincRNAs yielded amplified products of appropriate size as described in **Supplementary document Table 4**, while the negative control of DNase-treated RNA showed no amplification. The expression level of lincRNAs (by the band intensity) measured in RT-PCR were largely consistent with that measured by RNA-Seq for each line when the same amount of cDNA were used. Four lincRNAs (LIN11, LIN16, LIN27 and LIN32) showed additional bands besides the expected products, which could be alternative splicing transcript variants (or isoforms) of the same gene. While majority of selected lincRNAs, including eight NATs (LIN17 - LIN25) were found to express ubiquitously in both lines, six lincRNAs showed more line specific (LIN27 - LIN29 only expressed in line 216, whereas, LIN30 - LIN32 only expressed in line 218). LIN17-LIN25 were NATs showing evolutionary conservation against Arabidopsis TAIR10 intergenic sequences.

**a**

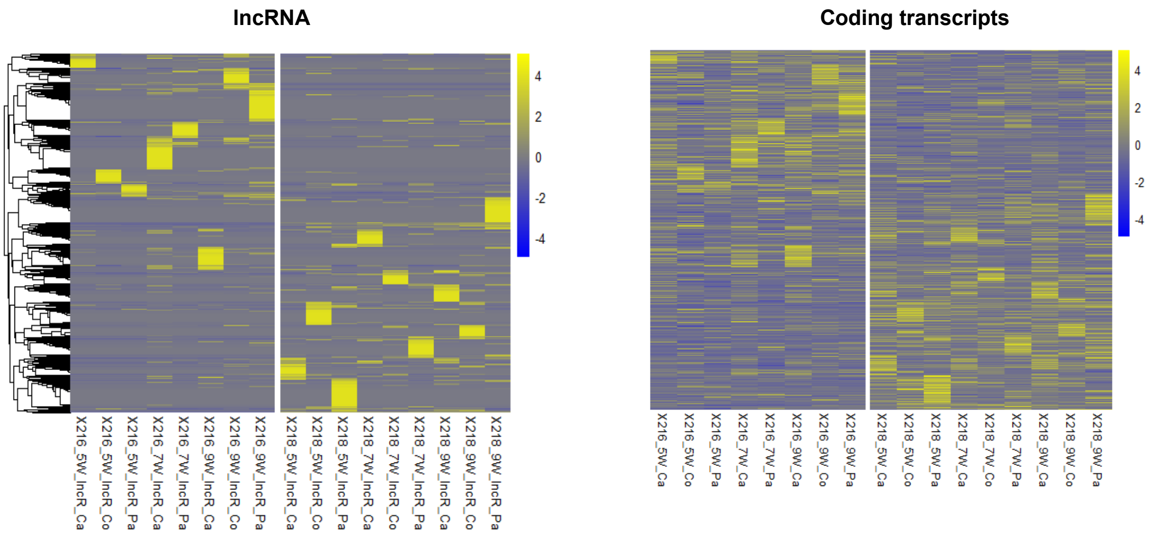


**b**

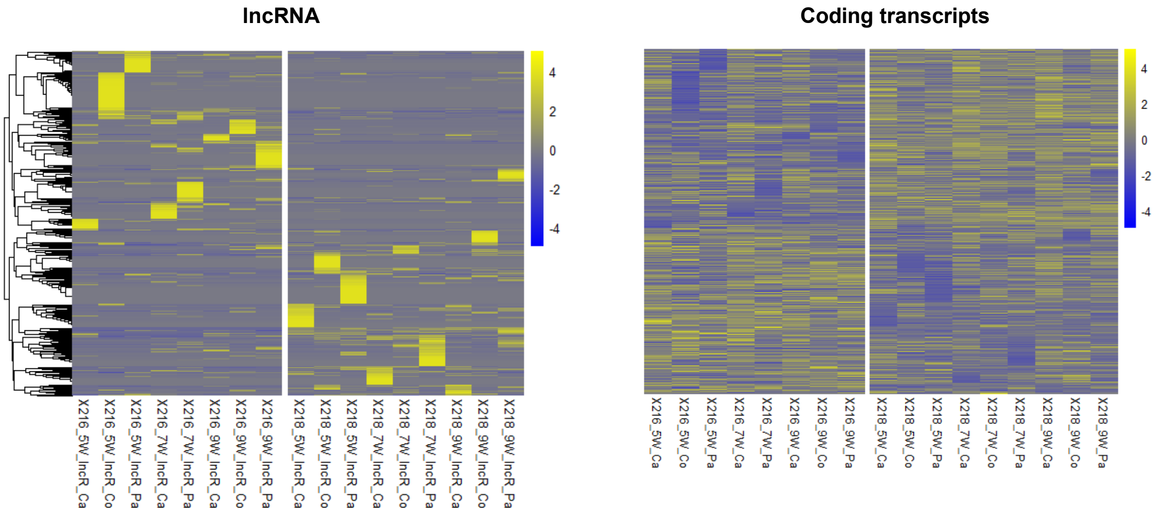


**Supplementary Figure 5. Heatmaps of expression of lncRNAs vs their associated coding transcripts. a** 1,533 pairs which showed positive correlation ($r \geq 0.3$, obtained from Pearson's correlation analysis). **b** 397 pairs of lincRNAs and neighboring genes which showed negative correlation ($r \leq -0.3$, obtained from Pearson's correlation analysis).

**a**

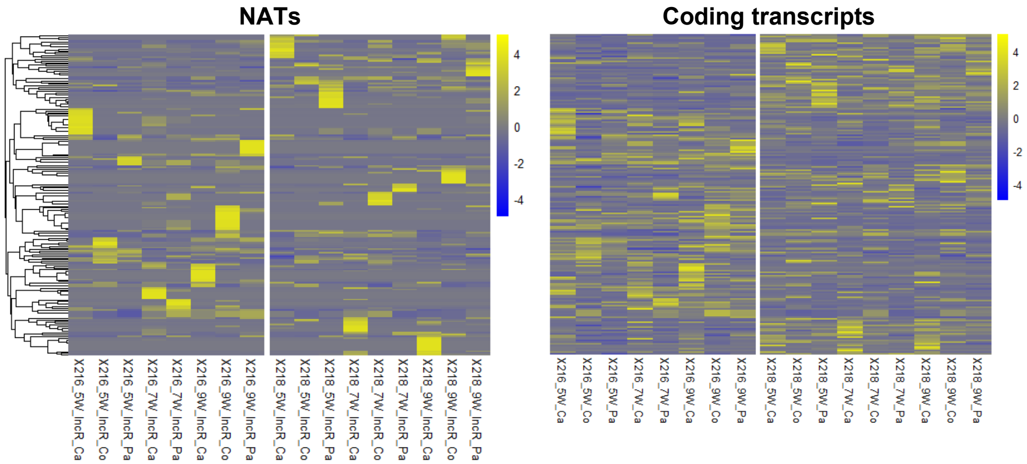


**b**

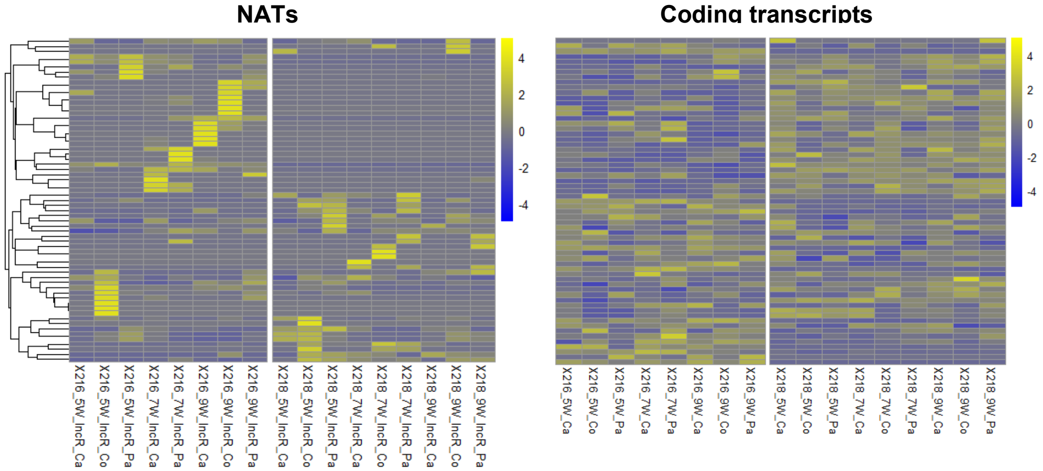


**Supplementary Figure 6. Heatmaps of expression of natural antisense transcripts (NATs) vs *trans* coding transcripts. a** 194 pairs of NAT-coding transcripts showing positive correlation ($r \geq 0.3$, obtained from Pearson's correlation analysis). **b** 68 pairs of NAT-coding transcripts showing positive correlation ($r \leq$-0.3, obtained from Pearson's correlation analysis).

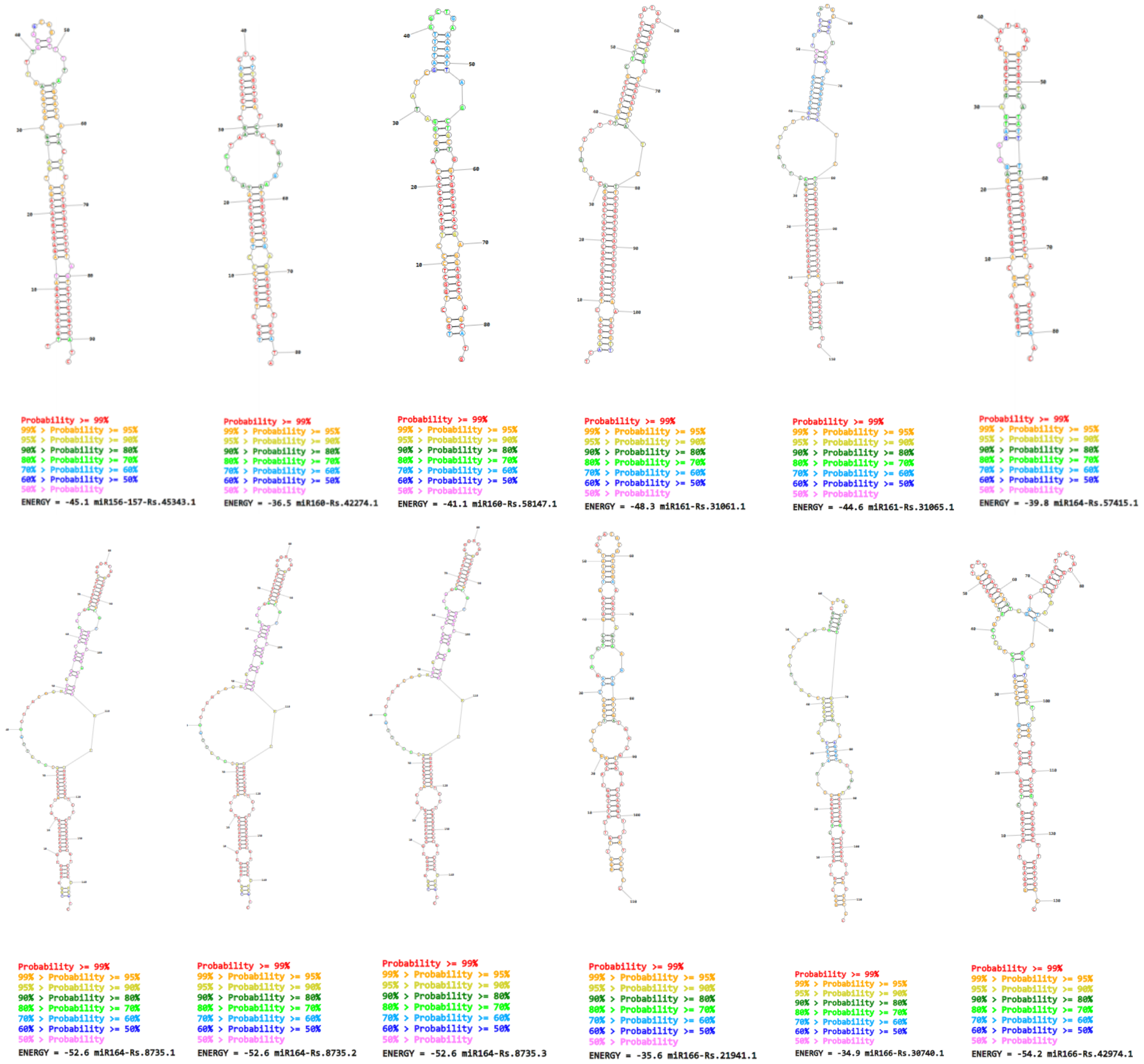


**Supplementary Figure 7. Secondary structure of 32 radish miRNA precursors identified in this study.** The secondary structures were predicted by RNAstructure web server with default settings (see Methods). The list of 32 radish miRNA precursors can be found in Supplementary Table Set 5 Table S3.

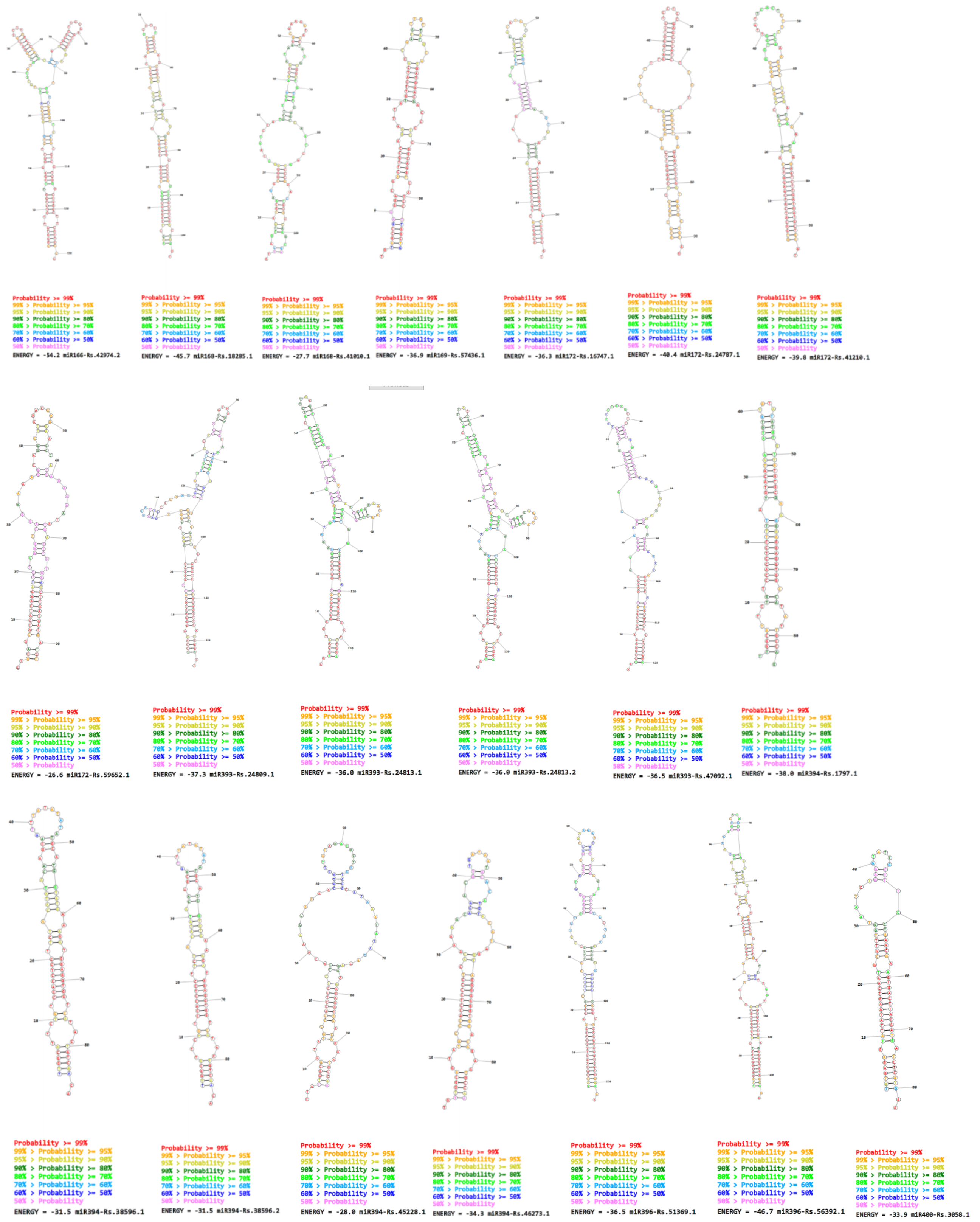


**Supplementary Figure 7.** Continued.

**a**

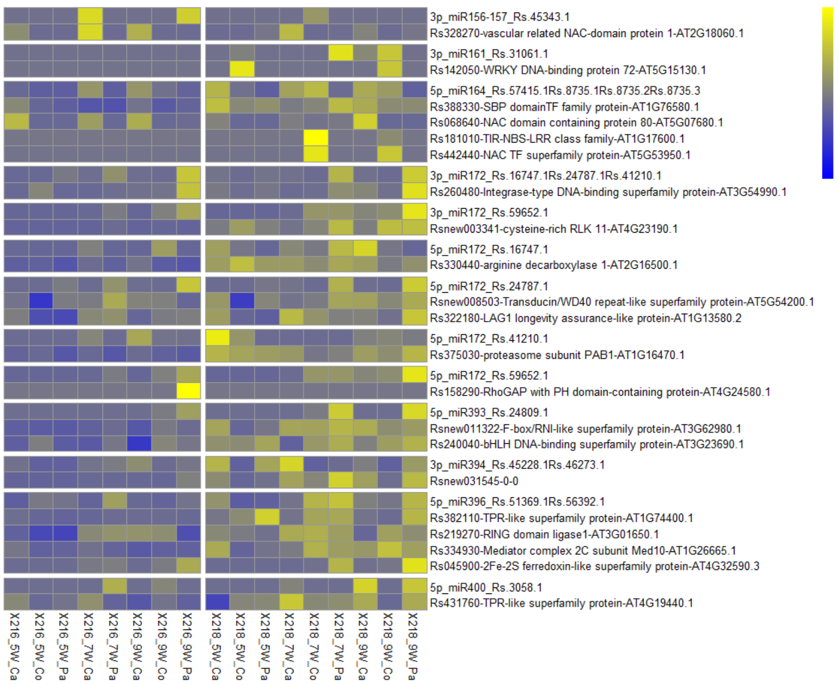


**b**

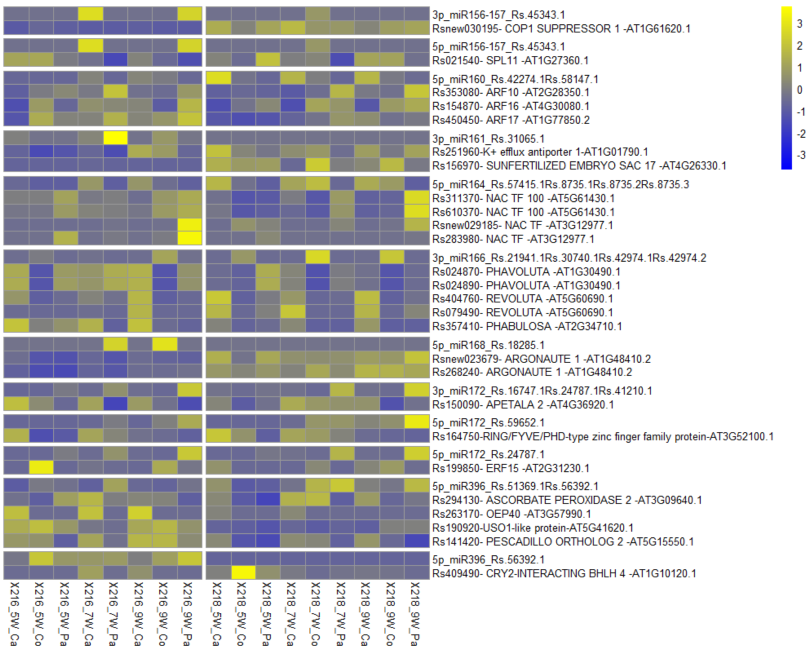


**Supplementary Figure 8. Heatmaps of expression of miRNA-encoding lincRNAs and target genes.** Target genes of miRNAs were identified by using the plant small RNA target analysis server - psRNATarget ver2017 (see Methods). **a** 26 pairs of miRNA precursors - target coding genes showing positive correlation ($r \geq 0.3$, obtained from Pearson's correlation analysis). **b** 36 pairs of miRNA precursors - target coding genes showing positive correlation ($r \leq$-0.3, obtained from Pearson's correlation analysis).

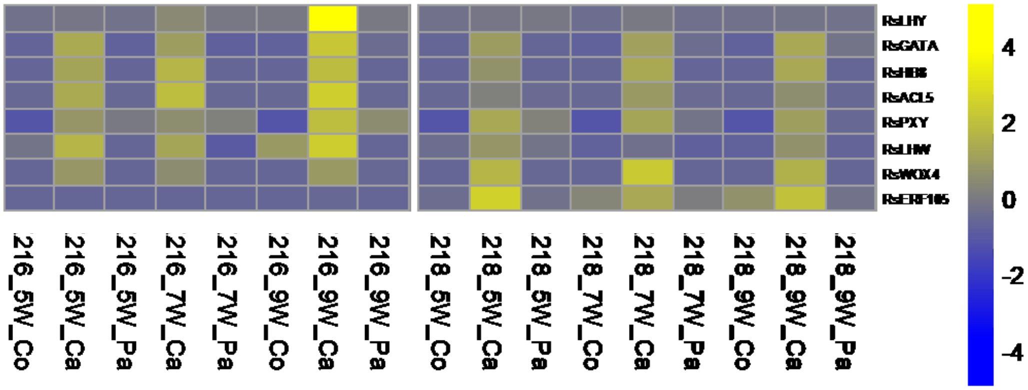


**Supplementary Figure 9. Row normalized expression of genes selected for RNA *in situ* hybridization.**

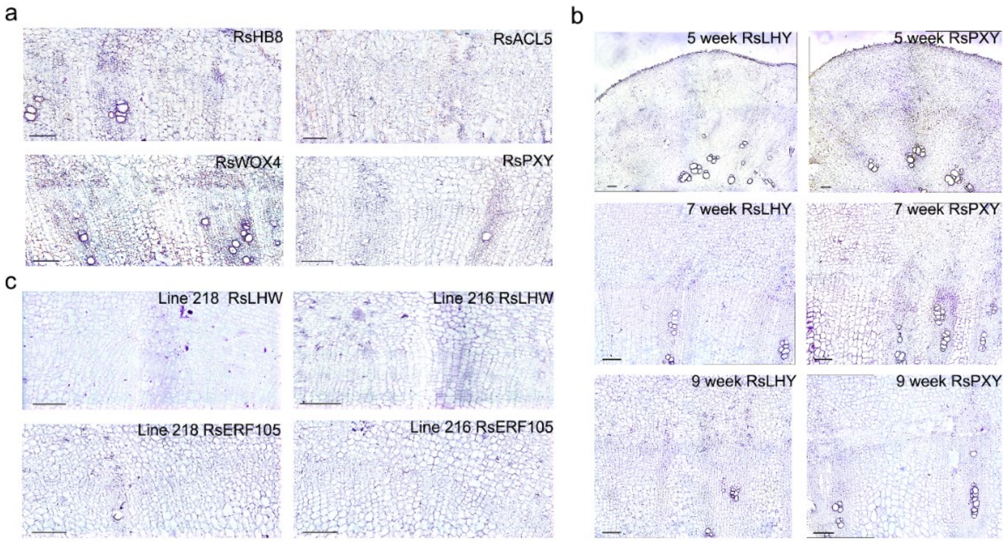


**Supplementary Figure 10. RNA *in situ* hybridization with sense probes. a-c** show *in situ* hybridization sense probe results. **a** *RsHB8*, *RsACL5*, *RsPXY* and *RsWOX4* in 7-week radishes of line 216. **b** shows a time course RNA-Seq validation in line 216 radishes of 5, 7 and 9 weeks using *RsPXY* and *RsLHY*. **c** shows cross-line comparison validation in 7-week radishes of line 216 and line 218 using *RsERF105* and *RsLHW*. . Magnification 100X, 200μm scale bar.

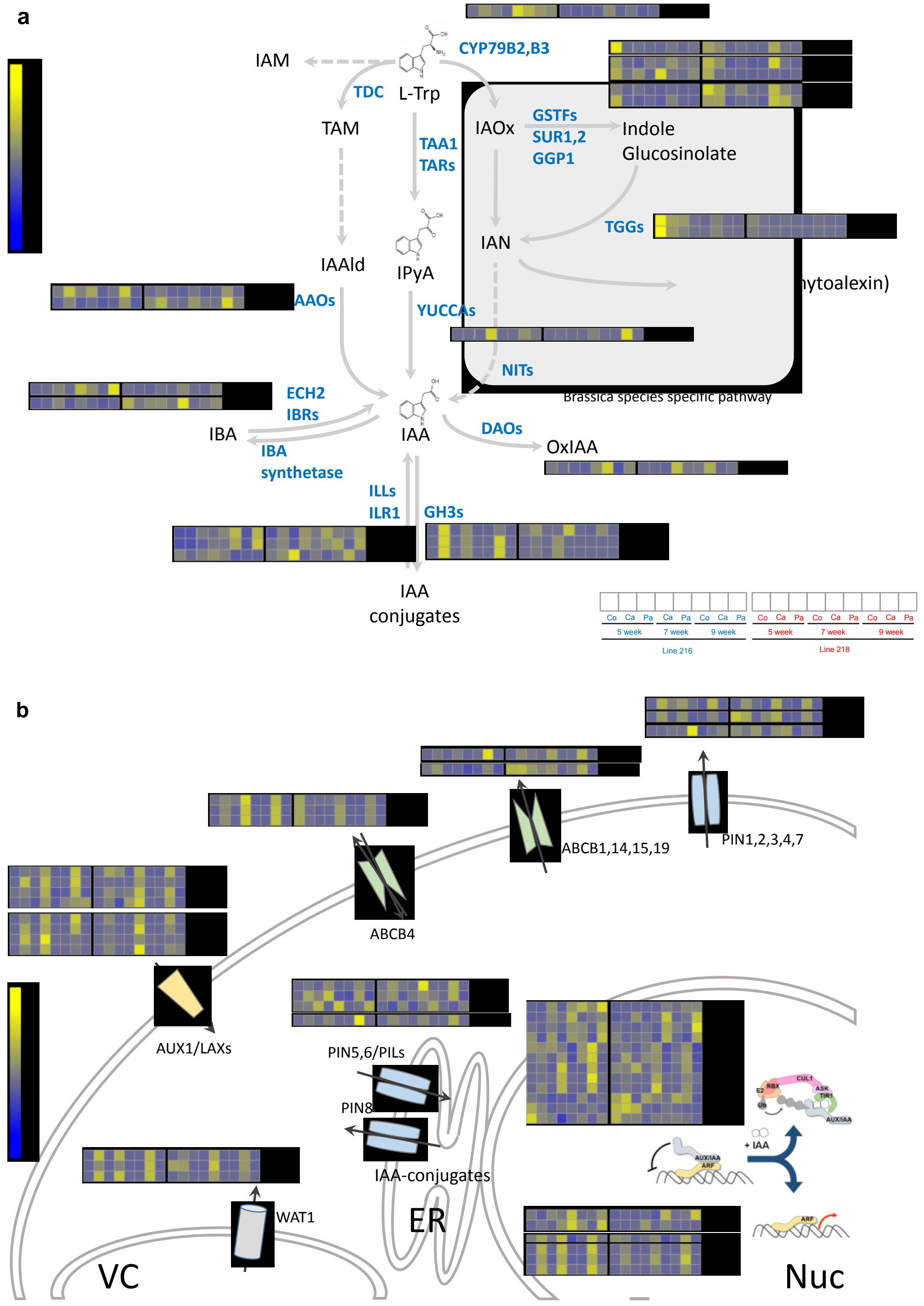


**Supplementary Figure 11. Auxin pathways. a** Schematic representation of known auxin biosynthesis pathway in Arabidopsis. Row normalized radish DEGs for the Arabidopsis genes in the pathway were visualized as heatmaps. **b** Schematic representation of known auxin transport and signaling in Arabidopsis. Row normalized radish DEGs for the Arabidopsis genes in the pathway were visualized as heatmaps. The order of sample was shown on the top right. Co, cortex; Ca, cambium, and Pa, parenchyma

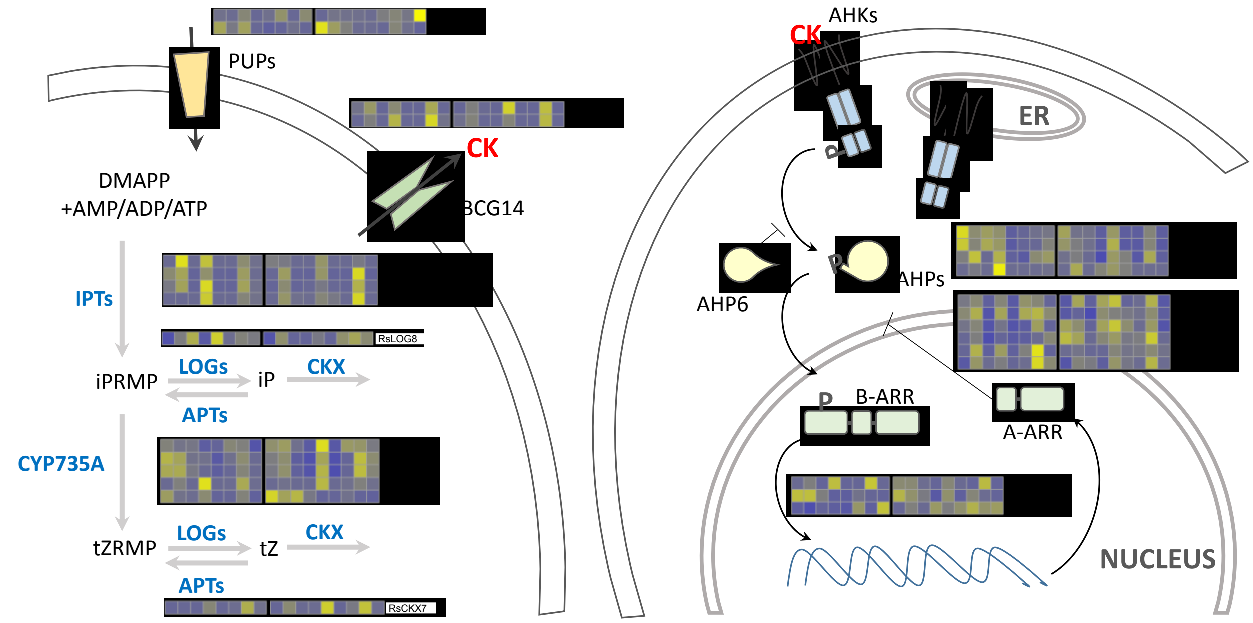


**Supplementary Figure 12. Cytokinin biosynthesis and signaling pathway.** Schematic representation of known cytokinin pathways in Arabidopsis. Row normalized radish DEGs for the Arabidopsis genes in the pathway were visualized as heatmaps. The order of sample was shown on the top right. Co, cortex; Ca, cambium, and Pa, parenchyma

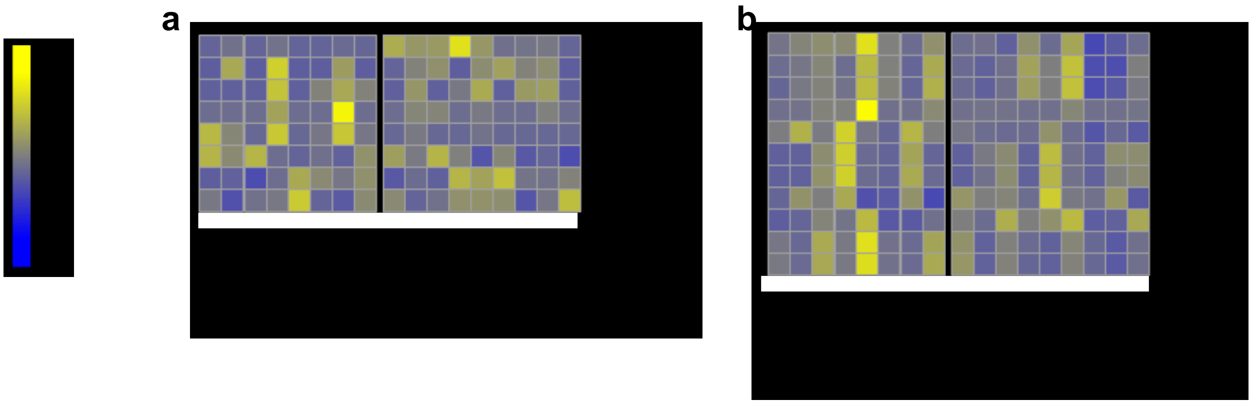


**Supplementary Figure 13. BR and GA pathway.** Row normalized radish DEGs for the Arabidopsis genes in the GA (a) and BR (b) pathways were visualized as heatmaps.

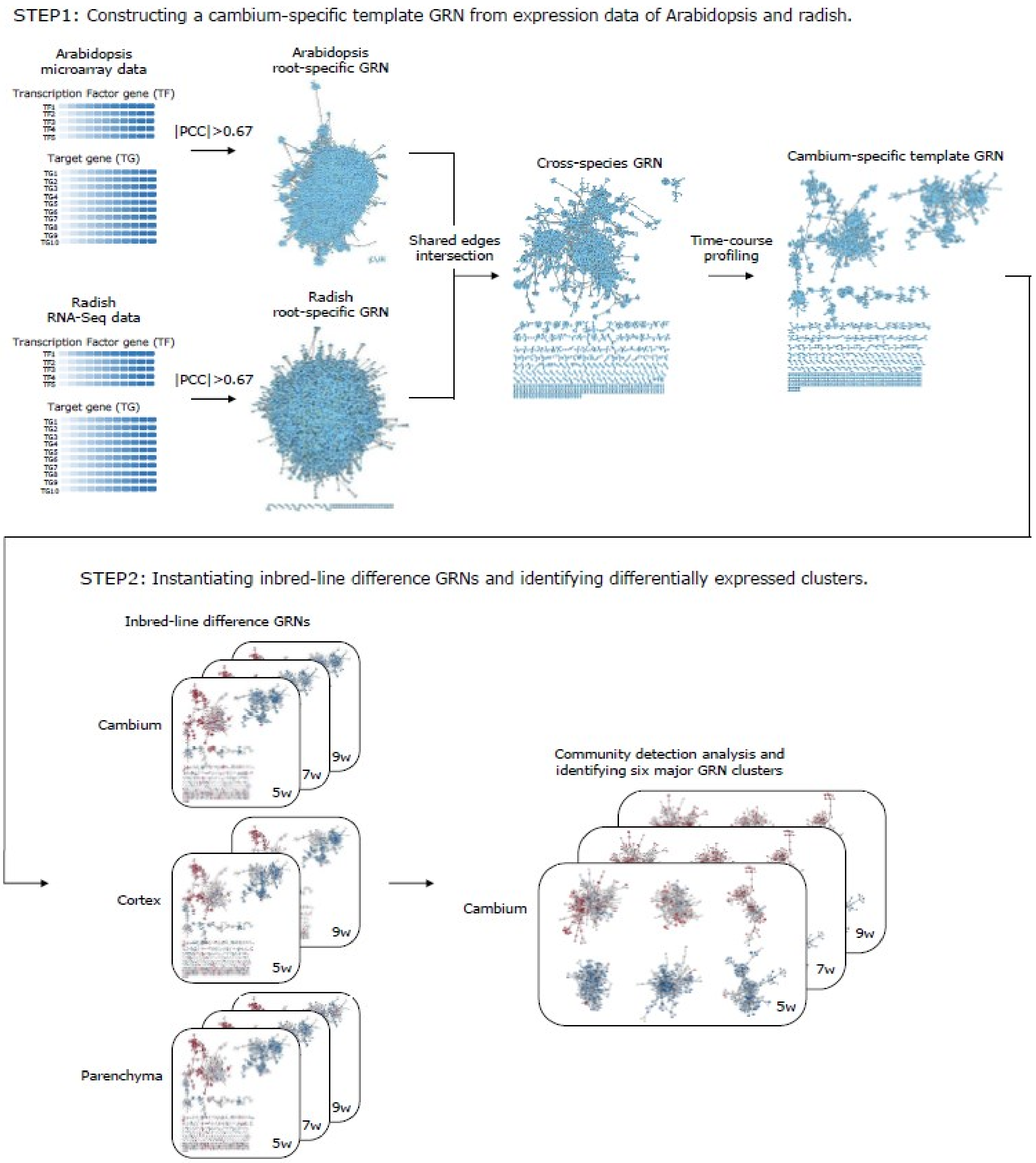


**Supplementary Figure 14. Workflow for Arabidopsis and radish cross-species network analysis.** The workflow consisted of two steps. In step 1, "Arabidopsis root-specific gene regulatory network (GRN)" and "radish root-specific GRN" were constructed by measuring all pairwise Pearson's correlation coefficients (PCCs) for TF-TF and TF-target gene pairs. A "cross-species GRN" was constructed by intersecting the shared edges of two networks. A "cambium-specific template GRN" was constructed by profiling cambium time-course transcription data for each gene. In step 2, "inbred-line differential GRN" was instantiated by mapping gene expression differences of inbred-lines 216 and 218 for each tissue type at each time point. Finally, a community detection analysis identified six major GRN clusters based on a statistical test with gene expression and GO terms.

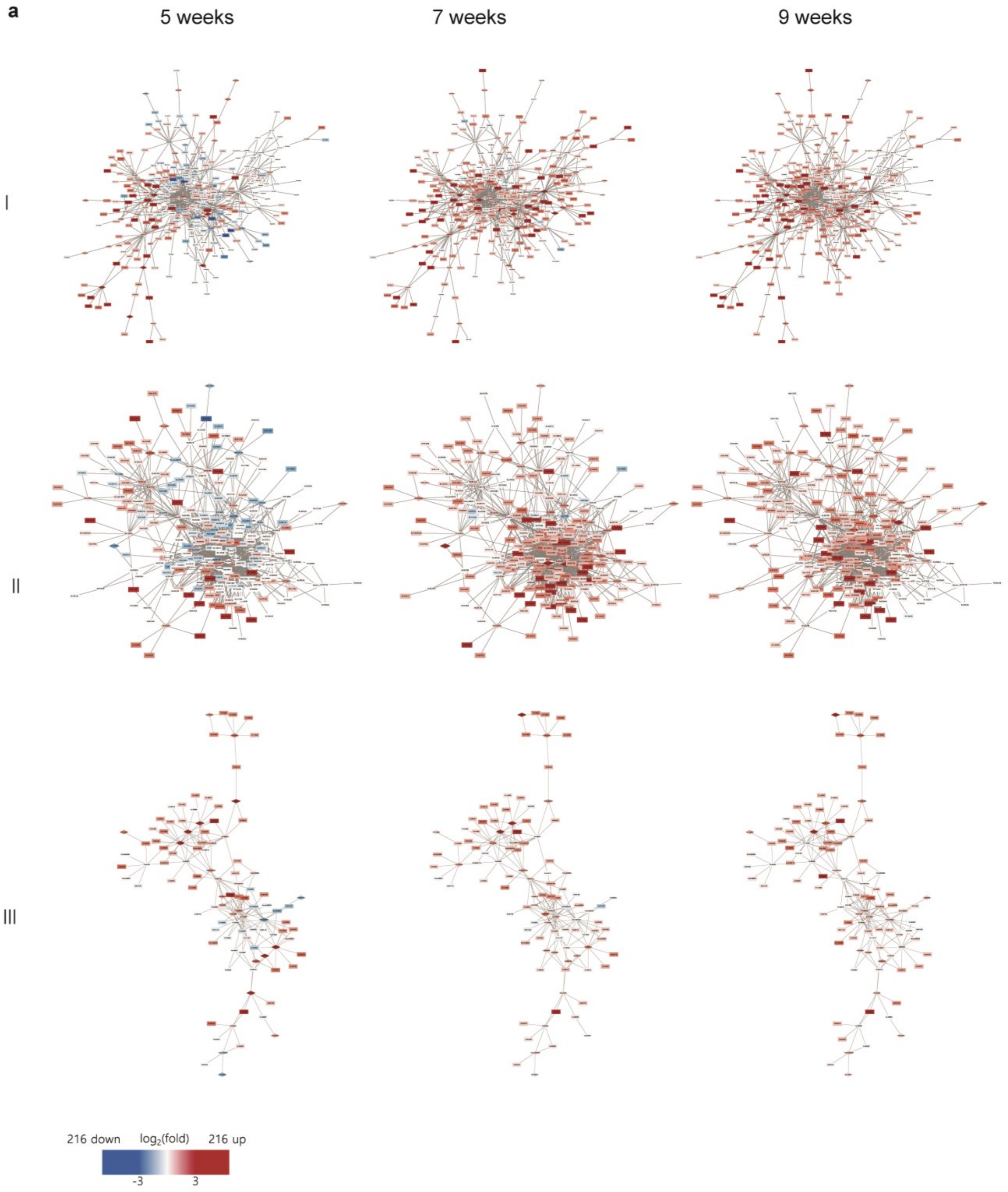
a
5 weeks
7 weeks
9 weeks
I
II
III
216 down
log2(fold)
216 up
-3
3

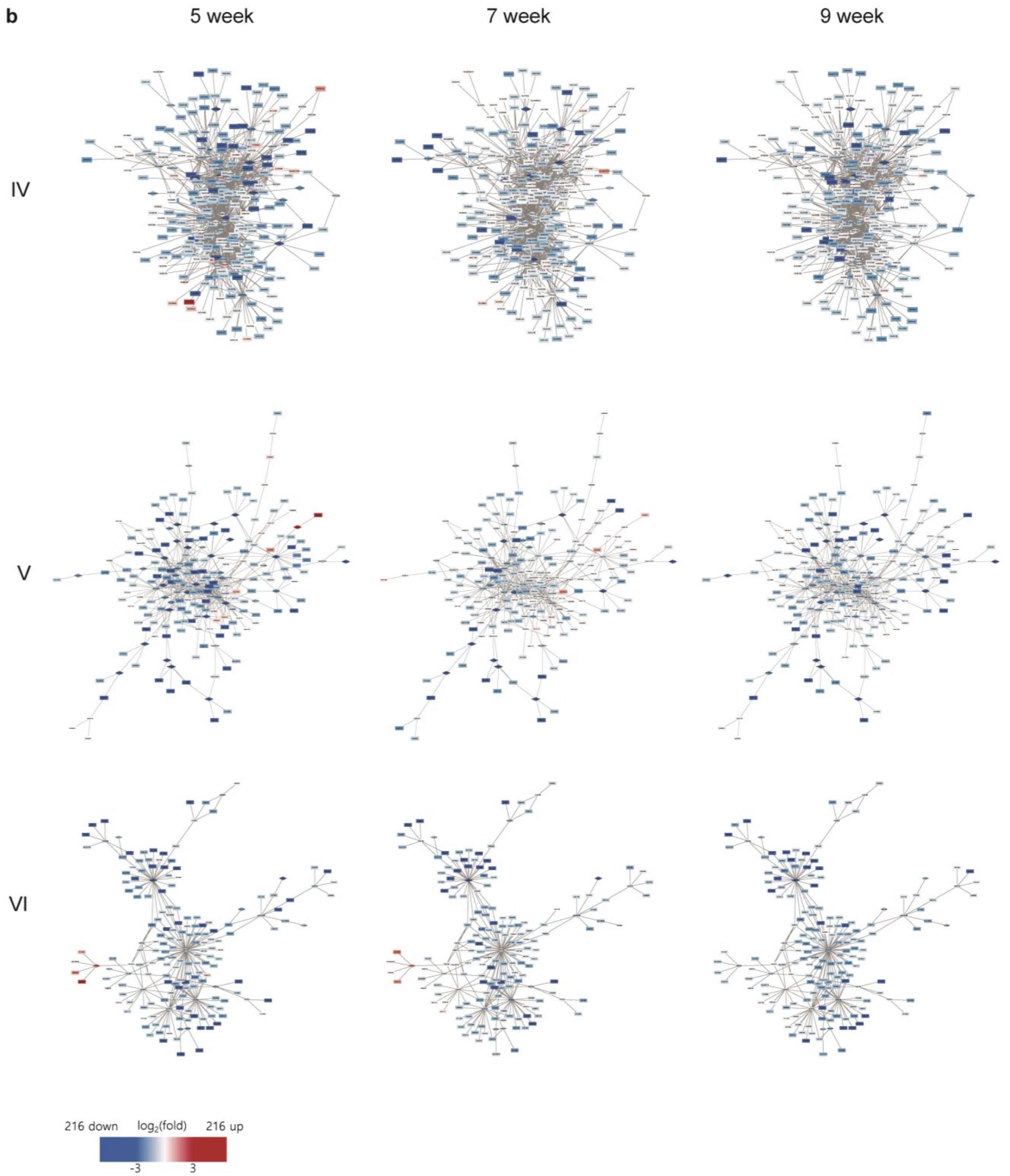


**Supplementary Figure 15. Conserved radish GRN clusters showing the relative expression of each node in the cambium in comparison between 216 and 218. a** Clusters showing the up-regulation in the cambia of line 216. **b** Clusters showing the up-regulation in the cambia of line 218.

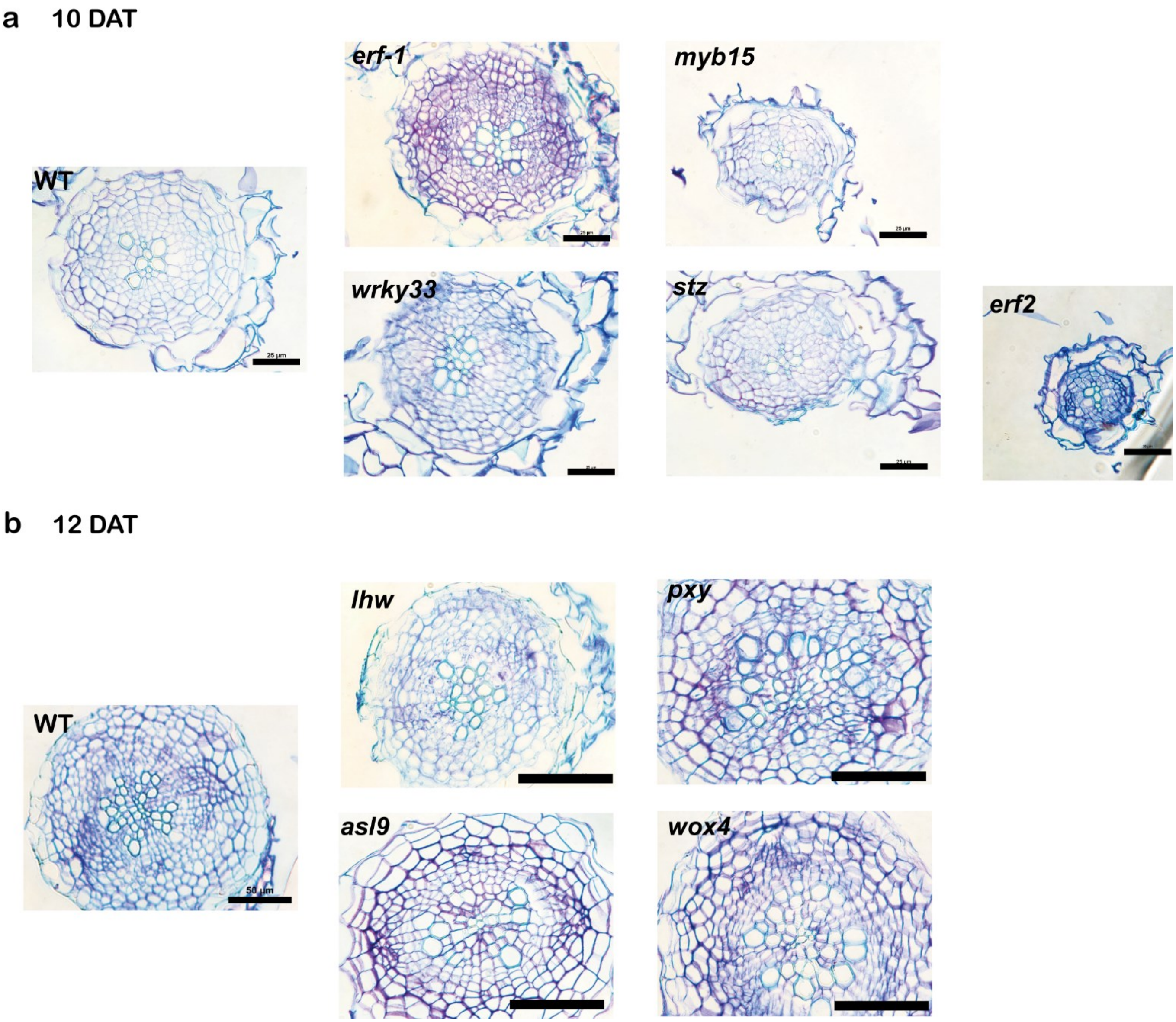


**Supplementary Figure 16. Root cross-section images for knockout mutant lines used in the *in vivo* GRN analyses. a** Cross section images of loss-of-function mutants of stress response transcription factor genes. Images were taken from 10 day old seedling roots. Scale bar, 25μm. **b** Cross section images of loss-of-function mutants of secondary growth regulators. Images were taken from 12 day old seedling roots. Scale bar, 50μm.

a

b

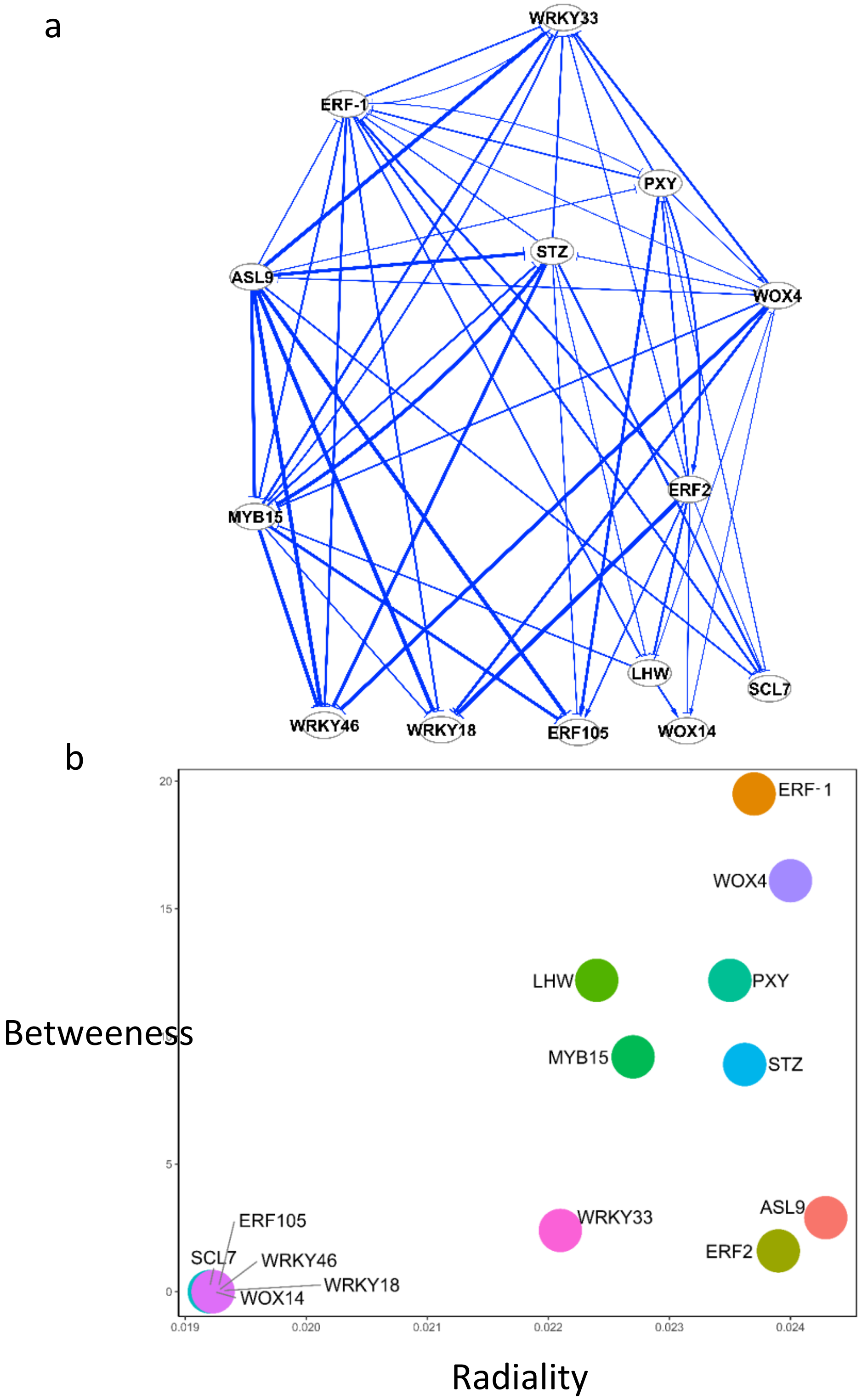


**Supplementary Figure 17.** GRN network (a) and scatter plot of radiality vs. betweenness (b) constructed from qRT-PCR data from the second batch of perturbation experiment

**Supplementary document references**


Ahn, H., Jung, I., Shin, S.J., Park, J., Rhee, S., Kim, J.K., Jung, W., Kwon, H.B., and Kim, S. (2017). Transcriptional Network Analysis Reveals Drought Resistance Mechanisms of AP2/ERF Transgenic Rice. Front Plant Sci 8:1044.

Axtell, M.J., and Meyers, B.C. (2018). Revisiting Criteria for Plant MicroRNA Annotation in the Era of Big Data. Plant Cell 30:272-284.

Dai, X., and Zhao, P.X. (2011). psRNATarget: a plant small RNA target analysis server. Nucleic Acids Res 39:W155-159.

Guo, L., and Lu, Z. (2010). The Fate of miRNA* Strand through Evolutionary Analysis: Implication for Degradation As Merely Carrier Strand or Potential Regulatory Molecule? PLOS ONE 5:e11387.

Huang, d.W., Sherman, B., and Lempicki, R. (2009). Systematic and integrative analysis of large gene lists using DAVID bioinformatics resources. Nat Protoc 4:44-57.

Jin, J., Tian, F., Yang, D.C., Meng, Y.Q., Kong, L., Luo, J., and Gao, G. (2017). PlantTFDB 4.0: toward a central hub for transcription factors and regulatory interactions in plants. Nucleic Acids Res 45:D1040-D1045.

Kozomara, A., and Griffiths-Jones, S. (2014). miRBase: annotating high confidence microRNAs using deep sequencing data. Nucleic Acids Res 42:D68-73.

Okamura, K., Phillips, M.D., Tyler, D.M., Duan, H., Chou, Y.T., and Lai, E.C. (2008). The regulatory activity of microRNA* species has substantial influence on microRNA and 3' UTR evolution. Nat Struct Mol Biol 15:354-363.

Pearson, K. (1895). Note on regression and inheritance in the case of two parents. Proc. R. Soc. Lond. B. Biol. Sci. 58:240-242.

Raices, M., Bukata, L., Sakuma, S., Borlido, J., Hernandez, L.S., Hart, D.O., and D'Angelo, M.A. (2017). Nuclear Pores Regulate Muscle Development and Maintenance by Assembling a Localized Mef2C Complex. Dev Cell 41:540-554 e547.

Reuter, J.S., and Mathews, D.H. (2010). RNAstructure: software for RNA secondary structure prediction and analysis. BMC Bioinformatics 11.

Su, G., Kuchinsky, A., Morris, J.H., States, D.J., and Meng, F. (2010). GLay: community structure analysis of biological networks. Bioinformatics 26:3135-3137.

Swarbreck, D., Wilks, C., Lamesch, P., Berardini, T.Z., Garcia-Hernandez, M., Foerster, H., Li, D., Meyer, T., Muller, R., Ploetz, L., et al. (2008). The Arabidopsis Information Resource (TAIR): gene structure and function annotation. Nucleic Acids Res 36:D1009-1014.

Untergasser, A., Cutcutache, I., Koressaar, T., Ye, J., Faircloth, B.C., Remm, M., and Rozen, S.G. (2012). Primer3--new capabilities and interfaces. Nucleic Acids Res 40:e115.

Xu, Y., Zhu, X., Gong, Y., Xu, L., Wang, Y., and Liu, L. (2012). Evaluation of reference genes for gene expression studies in radish (Raphanus sativus L.) using quantitative real-time PCR. Biochem. Biophys. Res. Commun. 424:398-403.

Yoder, A.C., Guo, K., Dillon, S.M., Phang, T., Lee, E.J., Harper, M.S., Helm, K., Kappes, J.C., Ochsenbauer, C., McCarter, M.D., et al. (2017). The transcriptome of HIV-1 infected intestinal CD4+ T cells exposed to enteric bacteria. PLoS Pathog 13:e1006226.

Yu, D., Meng, Y., Zuo, Z., Xue, J., and Wang, H. (2016). NATpipe: an integrative pipeline for systematical discovery of natural antisense transcripts (NATs) and phase-distributed nat-siRNAs from de novo assembled transcriptomes. Scientific reports 6:21666.

Zhang, J., Eswaran, G., Alonso-Serra, J., Kucukoglu, M., Xiang, J., Yang, W., Elo, A., Nieminen, K., Damén, T., Joung, J.-G., et al. (2019). Transcriptional regulatory framework for vascular cambium development in Arabidopsis roots. Nature Plants 5:1033-1042.
